# Ecological speciation in European whitefish is driven by a large-gaped predator

**DOI:** 10.1101/543744

**Authors:** Gunnar Öhlund, Mats Bodin, Karin A. Nilsson, Sven-Ola Öhlund, Kenyon B. Mobley, Alan G. Hudson, Mikael Peedu, Åke Brännström, Pia Bartels, Kim Præbel, Catherine L. Hein, Petter Johansson, Göran Englund

## Abstract

Lake-dwelling fish that form species pairs/flocks characterized by body size divergence are important model systems for speciation research. While several sources of divergent selection have been identified in these systems, their importance for driving the speciation process remains elusive. A major problem is that in retrospect, we cannot distinguish selection pressures that initiated divergence from those acting later in the process. To address this issue, we reconstructed the initial stages of speciation in European whitefish (*Coregonus lavaretus*) using data from 357 populations of varying age (26-10 000 years). We find that whitefish speciation is driven by a large-growing predator, the northern pike (*Esox lucius*). Pike initiates divergence by causing a largely plastic differentiation into benthic giants and pelagic dwarfs; ecotypes that will subsequently develop partial reproductive isolation and heritable differences in gill raker number. Using an eco-evolutionary model, we demonstrate how pike’s habitat specificity and large gape size are critical for imposing a between-habitat trade-off, causing prey to mature in a safer place or at a safer size. Thereby, we propose a novel mechanism for how predators may cause dwarf/giant speciation in lake-dwelling fish species.

## Introduction

For several decades, the question of whether speciation can occur in the face of homogenizing gene flow was hotly debated in evolutionary biology. Today, this debate has shifted focus as research has become more occupied with understanding the processes that cause speciation with gene flow in nature^1, 2^. Examples of on-going ecological speciation in sympatry are especially common in lake-dwelling fish, as they have an intriguing propensity to form genetically distinct ecotypes that differ in ecology, morphology, and reproductive biology^3, 4^. There is substantial variation among ecosystems and species as to how far this divergence has progressed, but a common feature is the evolution of large- and small growing ecotypes along resource- and/or habitat gradients in the lake environment. Examples of such ecotypic specialization include threespine sticklebacks^5^, African cichlids ^6^, rainbow smelt^7^, Arctic char^8^, Dolly Varden^9^, *Prosopium sp*.^10^ and a number of species belonging to the genus *Coregonus*^11–14^. Although the processes underlying this pattern have been studied intensively during recent decades^12, 15–19^, a fundamental question remains largely unanswered; why is divergence initiated in some populations and not in others?

To answer this question, we need to improve our understanding of how ecological mechanisms associated with habitat gradients could drive speciation. It is widely accepted that intense intraspecific competition and/or abundant ecological opportunities can cause divergent selection^20–27^. Other studies suggest that predation^6, 28, 29^, spatial variation in temperature^30^, environmental stress^31^, and reduced habitat- and prey availability^23^ can promote divergence. However, the importance of specific selective agents for actually causing speciation still remains elusive. A key problem is that divergence exposes incipient ecotypes to new ecological conditions, and the selective regime can change accordingly over time. For instance, if some ecological mechanism drives individuals to specialize in different habitats, this can cause divergent selection and conspicuous adaptations that are by-products rather than drivers of the initial divergence.

The best way to avoid confounding the causes and consequences of speciation is to study the process at its earliest stages^4, 32–39^. Unfortunately, this approach may lead us to study cases of early population divergence that are unrepresentative of the speciation process, or will never lead to speciation^4, 40^. This problem, in turn, could potentially be avoided by using comparative analyses^21, 23, 25, 26, 41, 42^ to identify the environmental conditions under which we can expect a future speciation process to proceed. So far, however, these two approaches have rarely been combined.

We addressed these issues by studying whitefish in Scandinavian lakes, where they form genetically distinct ecotype pairs that differ in body size^12^, morphology^12^, resource use^12, 43^, and time and place of spawning ^12^. Besides from being found in large numbers, these ecotype pairs are typically well known among local fishers^12^; opening up the possibility to use interviews as a method for collecting large amounts of spatial comparative data. Moreover, starting in the late 18^th^ century, there is a richly documented history of anthropogenic introductions that gave rise to new whitefish populations^44–46^. Today, the known and variable ages of these young populations provide an excellent opportunity to study how the speciation process initiates and develops over time. In this paper, we present extensive comparative data showing that northern pike is the key driver of ecological speciation in Scandinavian whitefish populations, and use data from populations of different age and modelling to form a hypothesis for why this large-growing predator is so critically important.

## Results

Our interviews with local fishers revealed that out of 357 lakes distributed from southern Norway to northern Sweden (Supplementary Fig. 1), 153 harboured ecotype pairs of dwarf and giant whitefish. These ecotype pairs were generally found in lakes with relatively high species-richness of fish, an observation that provides little support for the idea that intraspecific competition and ecological opportunity are the primary drivers of ecological speciation (Supplementary Fig. 2). Instead, our analyses showed that out of 13 analysed biotic and abiotic variables, presence of northern pike together with lake area and maximum depth determine divergence patterns of whitefish populations in our study area. Pike presence induces divergence into dwarfs and giants in lakes that are large and deep enough (Fig. 1, proportion correct predictions=0.90, Cohen’s κ=0.85). Smaller/shallower lakes have monomorphic whitefish, but the importance of pike for determining whitefish life histories can still be observed. Interview data on maximum weights showed that when pike are present, these lakes have either dwarf- or giant whitefish (Fig. 1a and Supplementary Fig. 3).

**Fig. 1:**
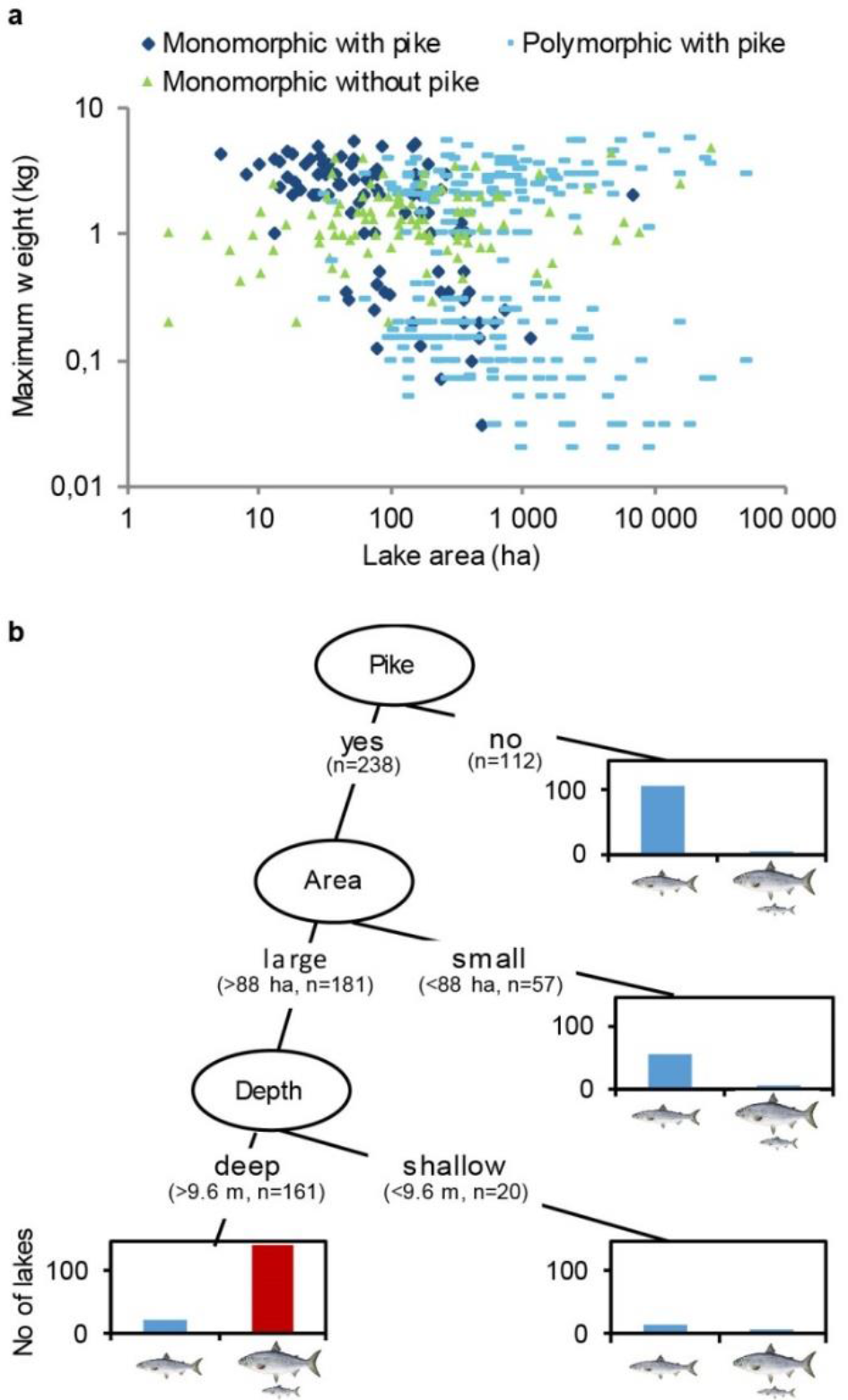
Pike presence, lake area and maximum depth control the formation of dwarf and giant whitefish ecotypes. **a)** Maximum weight (kg) of whitefish from populations in lakes with (n=217) and without (n=103) pike as a function of lake area. Light blue symbols represent polymorphic whitefish populations for which each lake has two corresponding observations. **b)** Classification tree (based on 13 explanatory variables, n=350) for the prevalence of polymorphism in whitefish, showing that pike induces co-occurring dwarf and giant ecotypes in lakes that are large and deep enough. The y-axes show the number of lakes. Cohen’s kappa for the whole model was 0.85.

Whenever fishers had knowledge about the spawning behaviour of whitefish in a lake with polymorphism, they reported segregation between dwarfs and giants in time and/or space during spawning. As dwarf- and giant whitefish ecotypes typically differ in a range of ecological traits^12^, this suggests that the dwarf/giant pairs reported in our interviews represent cases of incipient ecological speciation. To test this hypothesis and validate the fishers’ observations of polymorphism, we first looked for signs of reproductive isolation between sympatric dwarfs and giants using microsatellite data from 30 lakes where the interviewees had reported dual ecotypes. *F*_ST_-values were significant between dwarf and giant ecotypes in 23 of these lakes, and the non-significant differences between ecotypes were only found in young whitefish populations (introduced between 1825 and 1960, Supplementary Table 1). Second, we tested for differences in gill raker counts, a trait that is under strong genetic control in European whitefish (heritability *h*^2^ = 0.79 ^12, 47^), and known to be under divergent selection during resource specialization ^43, 48, 49^. When comparing dwarf and giant ecotypes in 70 lakes with reported size polymorphism, we found significant differences in 63 of them (Supplementary Table 2). Again, non-significant differences were only found in young populations. Restricting these analyses to populations that originated before the year 1900, we found that 21 out of 23 ecotype pairs had significant *F*_ST_-values (mean global *F*_ST_-value = 0.054, non-significant populations were introduced in Bölessjön, 1825 and Sörvikssjön, 1845, Supplementary Table 1), and that 61 out of 62 ecotype pairs differed significantly in gill raker counts (mean difference = 10.6 rakers, the non-significant one was introduced in Bomsjön, 1895, Supplementary Table 2). Except for in the youngest populations, our data thus show that the dwarf and giant ecotypes that fishers report have developed partial reproductive isolation and substantial differences in an ecologically important, heritable trait.

To understand how divergence initiates, we performed standardized gillnet sampling in 38 lakes that have recently introduced whitefish populations (introduced between 1784 and 1985); 23 where pike are present and 15 where they are absent. In order to only include lakes that are suitable for a future speciation process (see Fig. 1b), these sampling efforts were restricted to lakes that are larger than 100 hectares and deeper than 15 meters. First, we scanned the resulting data for signs of initiating body size divergence, and found a strong, rapidly appearing pike effect (Fig. 2). Body size variation among adult whitefish was larger in lakes with pike than in lakes without pike, and increased with time passed since introduction (ANCOVA: pike, t=5.33, p<0.00001, time since introduction, t=2.18, p=0.036, N=38). Separate analyses of the two lake categories showed that size variation increased with time in pike lakes (slope±SE=0.084±0.039, t=2.16, p=0.042, N=23), but not in lakes where pike were absent (slope±SE=-0.0079±0.031, t=0.26, p=0.80, N=15).

**Fig. 2:**
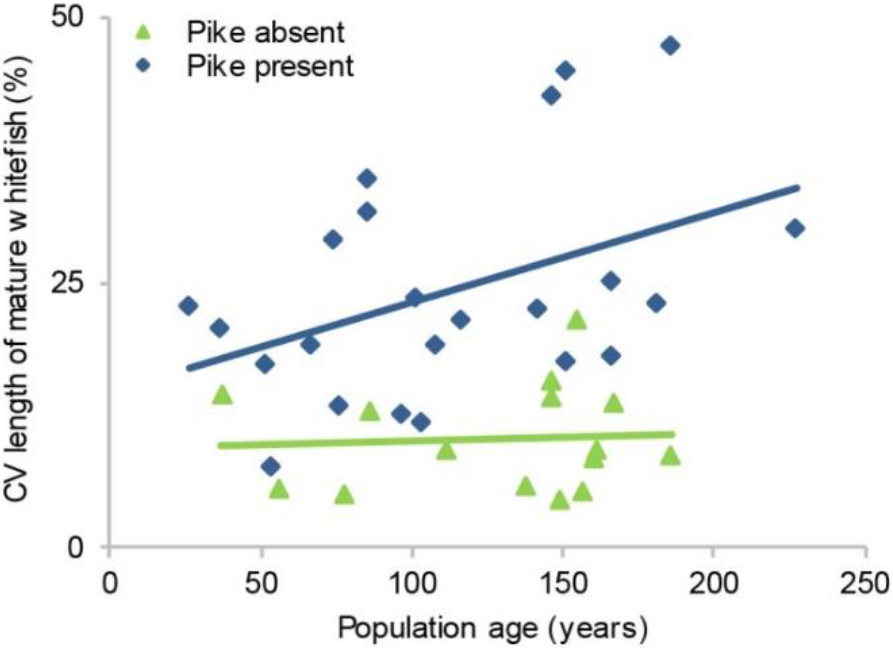
Pike presence drives rapid body size divergence in whitefish. Coefficient of variation for lengths of mature whitefish in lakes with (n=23) and without (n=15) pike as a function of population age.

Next, we used data from the sampled introduction-lakes to understand how this rapid body size divergence relates to divergence in other traits. Specifically, we compared the timing of divergence in body size and habitat use, which are highly plastic traits, with that of divergence in gill raker counts and neutral genetic markers. In these analyses, we wanted to exclude any pike-presence lakes where introductions of multiple genotypes may have contributed to the observed patterns of divergence. We therefore used microsatellite data (available for 18 lakes with known introduction dates) to exclude populations with signs of introductions of multiple genotypes, a procedure that left us with 11 populations with a putatively sympatric signal (Supplementary Fig. 4).

Going forward with these 11 populations, we first wanted to compare the initial divergence rates of body size and gill raker numbers. In order not to bias the comparison between the two traits, we performed cluster analyses ^50^ along the two trait axes simultaneously using the individuals caught in our standardized gillnet sampling. To allow comparison with much older populations, we also included samples from nine lakes with native, polymorphic whitefish.

The analyses gave divergent clusters in all populations except one (Lake Murusjøen, where whitefish were introduced in 1975). Analyzing how between-cluster differences in body size and gill rakers depend on population age (excluding Murusjøen), we found that divergence in body size is very rapid and precedes divergence in gill rakers (Fig 3, linear regression, divergence in body size: t=0.67, N=10, p=0.52; divergence in number of gill rakers: t=3.53, N=10, p=0.0076, both regressions excluding native populations). In fact, a large portion of the body size divergence typically seen in native polymorphic populations is expressed within just a few decades (Fig. 3). Moreover, a comparison between benthic and pelagic catches in the underlying gill net samples showed that this early size divergence is accompanied by an equally rapid divergence in habitat use between dwarfs and giants (Fig 4, ANCOVA excluding native lakes: pike, t=6.39, p<0.00001, time since introduction, t=1.17, p=0.26, N=20). Gill rakers on the other hand show very little divergence between the youngest clusters, suggesting that differences in body size form the basis for the initial formation of ecotypes (Fig 3).

**Fig. 3:**
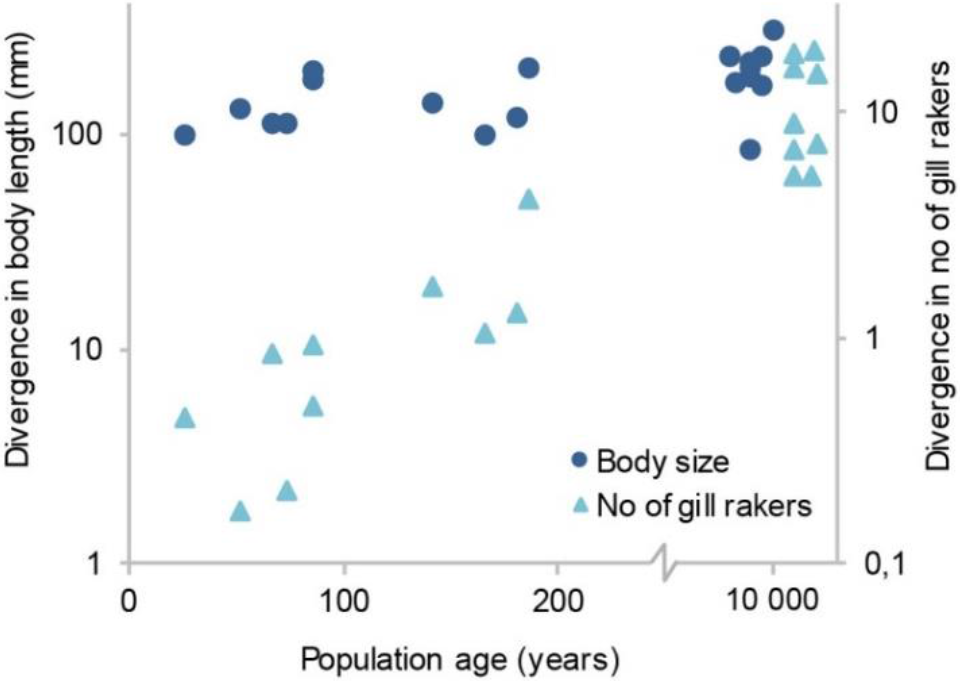
Rapid body size divergence leads the way to gill raker divergence. Between-cluster differences (based on mature individuals caught in our standardized gillnet surveys, n=19) in average values of body length and gill raker number as a function of population age. The positions of native populations were adjusted along the x-axis to reduce overlap.

**Fig. 4:**
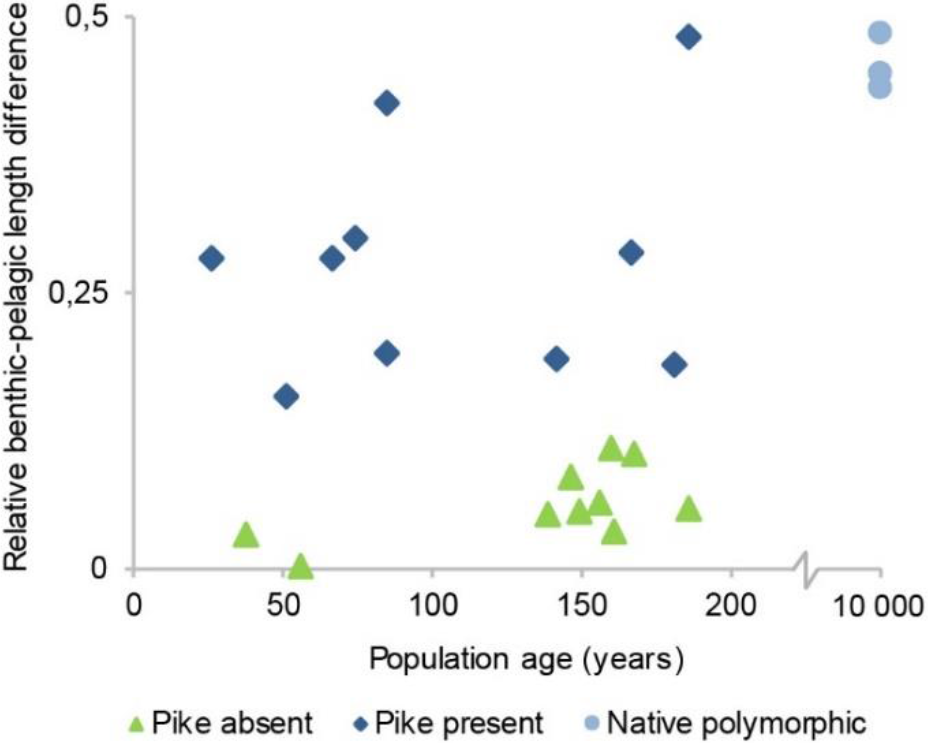
Body size divergence is associated with formation of benthic and pelagic ecotypes. Average length differences of sexually mature whitefish caught in littoral-benthic and pelagic gillnets as a function of population age. The figure includes the same selection of pike lakes that is presented in Fig. 3 (n=12, see methods section for information about missing data points) and pike-free control lakes (n=10). Length differences were calculated as (mean littoral length-mean pelagic length)/mean littoral length.

Next, we tested if gill raker counts differed between ecotypes within the 10 introduction lakes presented in Fig. 3. As the number of gill rakers could not be used for ecotype assignment in these tests, we classified individuals using body size and spawning site. The results from these analyses were consistent with the pattern resulting from our between-cluster comparisons. While gill raker numbers did not differ between dwarfs and giants in the youngest populations (introduced after year 1900, N=6, all t-values<1.68, all p>0.09, Supplementary Table 2), we found small but significant differences (1.5−2.7 rakers) in 4 out of 4 dwarf/giant pairs in the populations that were introduced during the 1800s (N=4, all t-values>3.22, all p<0.0017, Supplementary Table 2). The microsatellite data from these lakes showed a similar pattern; no significant population differentiation between ecotypes in the youngest populations but significant *F*_ST_-values between ecotypes in 2 out of the 4 older ones (Supplementary Table 1). Hence, the chronosequence of introduced populations suggests a timeline of divergence where the initial formation of dwarf/giant ecotypes is followed by more slowly appearing differences in gill raker numbers and neutral genetic markers (Figs. 3 and 4, Supplementary Tables 1 and 2).

Adding the native populations to the chronosequence, the short-term pike-driven divergence observed in introduced populations and the long-term pike-driven speciation process appear to form a continuum (Figs. 3 and 4). This suggests that we can view divergence in the youngest populations as representing the initial stages of the speciation process. Alternatively, it could be argued that size and habitat divergence may not necessarily lead to heritable differences and reproductive isolation, as has been observed in other fish species that form ecotypes^51, 52^. However, this does not appear to be the case in our study system. Surveying data from large and deep pike lakes with native whitefish, we did not find a single example of a dwarf/giant pair for which divergence had remained restricted to size (significant gill raker differences in 50/50 lakes, average difference =11.7; significant *F*_ST_-values in 13/13 lakes, average *F*_ST_=0.061). Hence, even though we lack direct experimental evidence, our data suggest that the initially formed dwarf and giant ecotypes with high predictability will continue to diverge along the speciation continuum.

The hypothesis that size differences lead the way to reproductive isolation implies that the spawning habits of whitefish will depend on their body size. In order to assess the validity of this corollary, we collected information (interview data validated with various kinds of sample fishing, see methods section and Supplementary Fig. 5 for details) about the average size of sexually mature individuals in all whitefish populations from our study lakes for which data on both spawning habitat and gill raker numbers were available. The resulting data showed that populations of giants typically spawn in shallow lake habitat, whereas more small-growing populations spawn either in streams, or in deeper water in the lakes (Fig 5). An analysis of this data confirmed that choice of spawning habitat is related to body size but not to gill raker counts (Multinomial logistic regression with stream spawners as reference; body size: stream vs shallow, Z=3.79, p=0.00015, stream vs deep, Z=2.13, p=0.033; gill rakers: stream vs shallow, Z=0.63, p=0.53, stream vs deep, Z=0.0025, p=1.0, N=72, Fig 5). All our empirical results thus point in the same direction; that the strong pike effect on whitefish divergence comes from a unique ability to induce pelagic dwarfs and benthic giants. To understand why pike have this ability as opposed to other potential predators (e.g. brown trout (*Salmo trutta*), arctic char (*Salvelinus alpinus*), and perch (*Perca fluviatilis*)), we must understand 1) how predation and it’s feedbacks on resource competition among prey can drive divergence into pelagic dwarfs and benthic giants, and 2) how this process depends on the characteristics of the focal predator species. Pike are largely restricted to the littoral zone of lakes and stands out by having a gape size large enough to catch relatively large prey ^53, 54^. To explore the consequences of the presence of a predator with these characteristics, we developed a size-structured eco-evolutionary model of the pike–whitefish system with whitefish maturation size as the evolving trait (see Supplementary Methods, Supplementary Figs. 6 and 7 and Supplementary Tables 3-5 for a detailed model description). We focused on maturation size because it is an important determinant of growth trajectories ^55^ that typically differs between sympatric ecotypes in our study system ^12^.

**Fig. 5:**
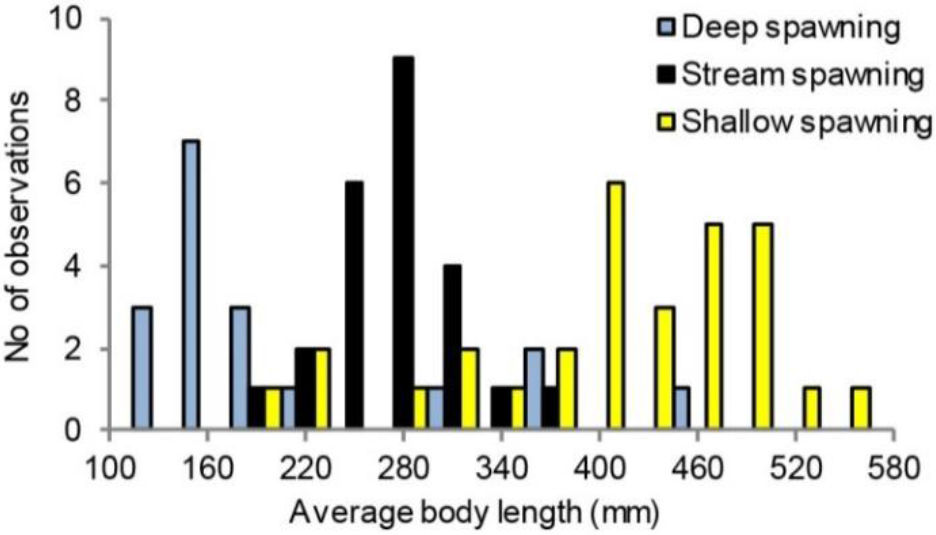
Whitefish spawning behaviour is related to body size. Histogram showing the distribution of average body lengths for populations that spawn in stream habitat, shallow lake habitat (depth <4 m), or deep lake habitat (depth >4 m) (n=72).

The model analyses suggest that habitat-specific predation can induce evolutionary divergence into dwarfs and giants by imposing a trade-off that affects life history and habitat choice of prey (Fig. 6a). The presence of pike causes whitefish to either 1) avoid pike in space at the cost of feeding on small pelagic zooplankton that provide limited scope for continued growth^56–58^, or 2) grow rapidly to reach a size that is subject to low predation risk by delaying the energy-consuming maturation and using the profitable littoral resource of large benthic invertebrates (Fig. 6c and Fig. 6d). A small-gaped predator does not impose this kind of trade-off (Fig. 6b), a result that corresponds well with our empirical data showing no association between whitefish divergence and the presence of small-gaped predator species such as brown trout, arctic char and perch. The mechanism behind the strong gape size effect is that when predation risk in the littoral habitat is confined to small prey, pelagic whitefish will be able to reach a size that allows them to shift to the littoral habitat without exposing themselves to high predation risk. Thus, two prerequisites for the necessary life history trade-off are 1) that the predator is sufficiently large-gaped to limit the ability of prey to grow out of the predation window when residing in the refuge habitat only, and 2) that prey can potentially reach sexual maturity before obtaining a safe size. Hence, besides prey–resource dynamics, the scope for this kind of predator-induced divergence will depend on a balance between the gape size of the predator and the inherent growth potential and life history of the prey.

**Fig. 6:**
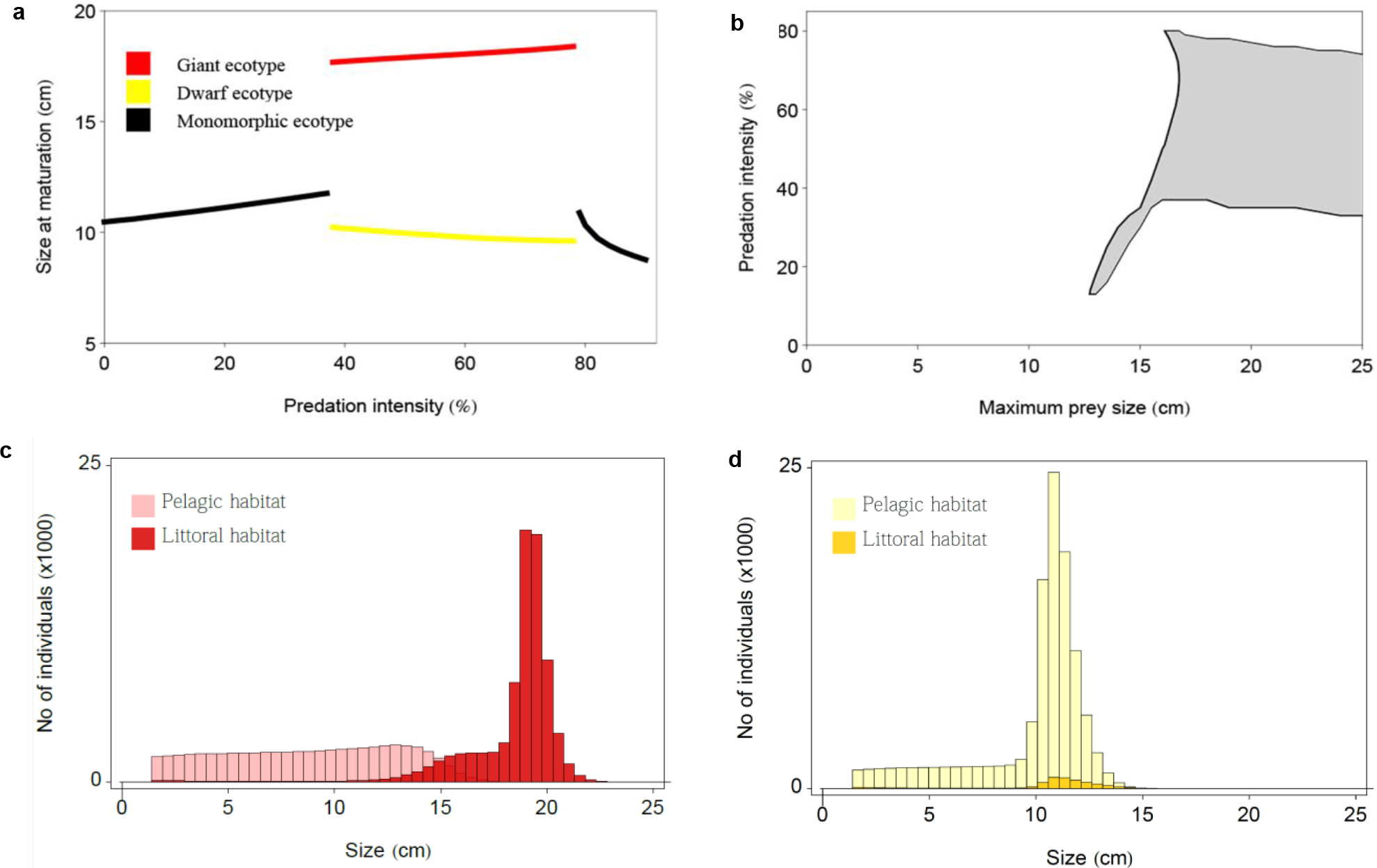
Large-gaped predators can induce dwarf- and giant prey ecotypes by imposing a habitat choice-growth strategy trade-off. a) Model simulation of maturation size as a function of predation intensity from a littoral predator capable of taking prey up to a maximum size of 18 cm. The red line represents giants that mature in the littoral habitat and the yellow line represents dwarfs that mature in the pelagic zone. b) The range of predation intensities (see Supplementary Table 4 for details) that induce evolutionary divergence at different values of maximum size of prey that can be taken by the predator. c) The distribution of the giant ecotype between the pelagic habitat and the littoral habitat at the evolutionary stable state (ESS) when the littoral predator can take prey up to 18 cm and the predation intensity is 70%. The giants mature at 18.2 cm. d) The corresponding distribution of the dwarf ecotype between the two habitats. Dwarfs mature at 9.7 cm.

## Discussion

In this study, we find an answer to the elusive question why benthic-pelagic ecotype pairs develop in some lakes and not in others. Contrary to popular belief, our data shows that ecological speciation along the benthic-pelagic habitat gradient is driven by a large-gaped predator. Recognizing pike’s critical role in our study system, we could then target the youngest pike-exposed whitefish populations to study the initial sequence of trait changes, and use a model rich in the necessary type of ecological detail to analyse the underlying mechanisms. The results suggest that pike drives ecological speciation by inducing pelagic dwarfs and benthic giants; a primary ecotypic differentiation that forms the basis for further divergent adaptations to the respective habitats, and at the same time promotes reproductive isolation.

While previous work has shown that both gill raker numbers and body size are under divergent selection during whitefish radiations^49^, our data thus suggest that divergent selection on body size and habitat use is the primary route to ecotypic differentiation and subsequent ecological speciation. While body size divergence has been described as an important component of niche differentiation during ecological speciation in other systems^6, 59^, the full, ecological implications of size differences have received relatively little attention in studies of ecological speciation in fish. Unlike other morphological traits, body size determines both an individual’s potential gain from feeding on a given food type and its exposure to predation risk while doing so^60^. As a consequence, small and large individuals that face between-habitat variation in resource gain and predation risk will often specialize on feeding in different habitats^61, 62^. At the same time, individual growth depends on the density and quality of available resources^60^, and feeding on small- or large prey can affect ontogenetic growth trajectories differently^56, 63^. This fundamental property of body size, i.e. that it both determines and is affected by an individual’s ecological niche, is a critical component of the trade-off that gives body-size divergence in our model. Hence, our findings are consistent with the idea that phenotypic plasticity is important for speciation^51, 64–66.^

Through the plasticity of food-dependent growth, inherently small-growing individuals can be scared into sacrificing growth opportunities, while inherently large-growing individuals can gain access to resources that allow continued, rapid growth. This way, food-dependent growth can greatly enhance the adaptive significance of heritable body size variation. In our model, such variation is represented by differences in maturation size; a major source of growth trajectory variation among fish populations ^55^, and a typical feature of whitefish radiations ^67^. However, any trait variation that affects individual growth could potentially sort individuals along a gradient of size-dependent resource gain/predation risk. Our model should therefore be viewed as the most straightforward representation of a more general idea; that gape limited predation can cause individual prey to either stay in refuge habitat, or maximize growth to reach a safe size, depending on their inherent growth potential.

While our results improve our understanding of how benthic and pelagic ecotypes form, they offer more limited insight into how giants and dwarfs continue to diverge towards speciation; a process that requires assortative mating and some form of heritability that transfers the growth strategies and their spawning behaviour between generations. Predation risk could potentially explain the association between body size and choice of spawning sites in much the same way as with size-dependent habitat choice outside of the spawning season. This remains to be tested, but the size-spawning site association nevertheless provides a plausible explanation for why dwarfs and giants would develop reproductive isolation over time. Our study thus contributes to a growing body of evidence suggesting that differences in body size may be an important driver of reproductive isolation in polymorphic fish populations^68, 69^. When it comes to the inheritance of adult size, a specific mechanism remains to be demonstrated. It could come from genetically controlled differences in maturation time or size, but there are other possible mechanisms by which size differences could be transferred between generations. For example, size-dependent choice of spawning sites could feed back on the hatching time and early growth of offspring because the different spawning habitats have different temperature regimes^15, 70^. Moreover, dwarfs produce smaller eggs than giants^71^ and it has been shown that this impedes the initial growth of their offspring^72^. Demonstrating the mechanisms that cause reproductive isolation between dwarfs and giants will be an important challenge for future research.

The phenomenon that fish populations form sympatric, large- and small-growing ecotypes has been repeated in a large number of species, and along all major habitat axes in lakes^5, 8, 73–75^. If this parallelism is mirrored in the underlying mechanisms, our results suggest that predation is heavily underestimated as a driver of intraspecific fish diversity in lakes. While our results apply directly to divergence along the benthic-pelagic resource axis, the type of trade-off that gives divergence in our model could appear along any gradient where small prey fish take refuge in suboptimal growth conditions. Such growth conditions can come from spatial variation in a range of environmental variables, and do not necessarily depend on the presence of discrete, habitat-specific resource types. Hence, predator-induced trade-offs could potentially explain why dwarf- and giant ecotypes form also in situations where diet specialization is less pronounced or even absent^8, 76–78^.

To test the hypothesis that predator-induced growth strategies are generally important as a starting point of ecological speciation, we need to disentangle leaders and followers among the selection pressures and diverging traits that are involved when ecotype pairs form. Our study illustrates how this can be achieved by combining comparative and temporal data, as this can allow us to both identify crucial selection pressures and study their effects on populations over time. Applied to a variety of systems and including a wide range of study methods, this approach holds great promise to improve our understanding of how ecology initiates speciation with gene flow.

## Methods

We used data from 357 Scandinavian lakes distributed along a south– north gradient from southern Norway (58.99 N, 8.29 E) to northern Sweden (68.17 N, 21.97 E) (Supplementary Fig. 1).

### Data collection

#### Interviews

Local fishers often have detailed knowledge about the habits and spawning sites of whitefish ecotypes in Scandinavian lakes, and co-occurring ecotypes typically have distinct local names^12^. This allowed us to use interviews to assess large-scale patterns of maximum body size and the frequency of polymorphism. We asked local fishers (and other persons with relevant knowledge) if the whitefish in a given lake was indigenous or introduced, if there were one or more ecotypes, and for the maximum weight, spawning site and spawning time of each ecotype. Care was taken to follow the same interview protocol for all lakes. In order to estimate maximum size, we asked about the largest specimen caught in a given lake during the last 25-year period. We used maximum weight as a crude life history metric because fishers tend to remember this figure and because it effectively captures the divergence between dwarfs and giants. We defined polymorphism as the existence of two or more coexisting populations with different maximum sizes. When deciding whether or not whitefish populations were polymorphic, lakes were divided into the following four categories. 1) Fishers report two or more populations with different maximum sizes that use different spawning grounds and/or differ in spawning time. 2) Fishers report two ecotypes that differ in maximum size but could not provide information about spawning. 3) Fishers report indications of polymorphism, such as presence of both large and dwarfed spawners and size-related differences in parasite load, but feel uncertain if these represent different ecotypes. 4) Fishers report that, to the best of their knowledge, there is only one ecotype of whitefish. In the final data set, we defined lakes from categories 1 (n=105) and 2 (n=51) as being polymorphic and lakes from category 4 (n=197) as being monomorphic. Lakes in category 3 (n=22) were excluded, with the exception of four lakes where we had performed standardized sample fishing.

For a subset of lakes (N=72) used to analyze the association between spawning habitat and phenotype (average body size and gill raker number, see below for more details), we also asked fishers about the water depth at the spawning sites and the average size of spawning individuals.

#### Publications and official records

Data from publications and official records were mainly used to assess the age and origin of populations, and a large proportion of the records of year of introduction in our data set originate from Swedish and Norwegian reports that were published between 1797 and 2013^44, 46, 79, 80^. In cases where the time of introduction was given in a time-span of up to 20 years, we used the middle year. Published data were also included in analyses of differences in gill raker counts and neutral genetic markers (Supplementary Tables 1 and 2). Finally, for the analyses of phenotype-spawning habitat correlation, we used published information about spawning depth^12, 81–83^ (N=11) and average body size^76, 81, 83, 84^ (N=7) for populations where interviews did not provide this information.

#### Field sampling

To validate interview data and to catch fish for genetic and phenotypic analyses, we performed standardized gillnet sampling in 51 of the interview lakes, and some form of non-standardized sampling (gillnetting, hand netting or ice fishing) on spawning grounds in 22 of them (or their adjacent streams).

As a standard gillnet setup, we used 24 benthic gillnets (30×1.5 m; 8 of multimesh-type, 4 with panels of 33 mm and 12 with 45 mm mesh size knot to knot) and 8 floating gillnets (two of multimesh type (27×6 m) and six single-meshed nets (30×5m) with mesh sizes 12, 15, 20, 23, 30 and 38 mm). In a subset of the sampled lakes, the standard setup was extended to include two extra floating gillnets with mesh sizes of 33 and 45 mm. Including these mesh sizes in the pelagic setup allowed us to use the combined catches from multimesh, 33mm and 45 mm nets to directly compare the average size of sexually mature whitefish in the benthic and pelagic habitats respectively (Fig. 4). We have performed this extended sampling in 13 of the lakes included in Fig. 3 and in 10 pike-free control lakes. Hence, the data points in Fig. 3 that are missing in Fig. 4 (6/9 of the native populations) are missing because the gillnet setup used in these lakes did not allow the relevant between-habitat comparison of average body size.

#### Phenotypic data

The number of gill rakers on the first left gill arch were counted under a dissecting microscope. We present gill raker data from ecotype pairs in 72 lakes, out of which 50 had putatively native- and 22 had introduced whitefish populations. In 35 of these lakes, the gill raker counts were based on our own samples, and in the remaining 37, we used published data (Supplementary Table 2). In lakes with more than two ecotypes, we compared the gill raker count of the largest and the smallest ecotype. For the analysis relating average phenotype to spawning habitat (see below), we recorded gill raker means for 10 additional lakes where data were available for only one population. Body length and sexual maturity status were recorded in the field.

#### Analyses

Our interview-based data set contains data from 357 lakes, and all non-interview based data comes from subsets of these lakes. Populations of recent, monomorphic origin cannot be expected to be polymorphic, and may experience rapidly changing growth conditions. Therefore, we did not include whitefish populations introduced after 1960 (the most recent introduction year that has given rise to a polymorphic population according to our interviews) in Fig. 1, Supplementary Figs. 2 and 3, and the underlying analyses. For all other analyses, we used the maximum number of lakes that was applicable and for which we had relevant data. This means that the number of lakes included in different analyses vary, either because interviews did not result in complete data for all questions, or because non-interview data was not available for all lakes.

For statistical analyses, including linear regression, ANCOVA, and t-test, we scanned residual plots for heteroscedasticity, outliers, and model misspecification. When motivated, we used logarithmic or square root transformations to reduce heteroscedasticity and the influence of outlying observations. For logistic regression analyses, we scanned Pearson and deviance residuals for outliers. No outliers or signs of model misspecification were detected.

#### Ecological drivers of polymorphism

Relationships between environmental variables and the prevalence of polymorphism were modelled with classification trees, estimated and crossvalidated with the *rpart* module in R^85^. Thirteen variables were used as predictors: the number of fish species co-occurring with whitefish, lake area, maximum depth, altitude, temperature sum (total number of degree days above 6 °C), and presence/absence of the fish species pike, roach (*Rutilius rutilus*), grayling (*Thymallus thymallus*), burbot (*Lota lota*), Eurasian perch (*Perca fluvialitis*), arctic charr (*Salvelinus alpinus*), brown trout (*Salmo trutta*), and European minnow (*Phoxinus phoxinus*). Optimal tree depth was determined with cross validation and the agreement between data and model predictions was judged with Cohen’s κ-statistics^86^.

#### Divergence vs population age

To analyse the relationship between population age and the degree of divergence in body size and gill raker counts (Fig. 3), we first used the procedure mclustICL in the R module mclust^50^ to identify clusters based on body length and gill raker counts from mature whitefish caught in our standardized gillnet sampling. Missing data were imputed with the imputeData command in the mix package^87^. The difference between mean values for the clusters in a lake was then used as a measure of divergence in body size and number of gill rakers. If more than two clusters were identified, we excluded the intermediate ones.

When gill raker comparisons were made between individuals that were preassigned to ecotype, we compared mature small individuals (<25 cm) and large individuals (>35 cm) caught either on their spawning grounds (dwarf sample from six lakes) or from sampling not associated to spawning grounds (dwarf sample from four lakes and giant sample from all 10 lakes). In one lake (Stor-Skirsjön), whitefish rarely grow larger than 35 cm, and we therefore compared the mature dwarfs (average length 182 mm) to fish >275mm.

#### Phenotype-spawning habitat correlation

Our analysis of the correlation between whitefish phenotype and spawning habitat included populations for which we could get information about spawning habitat and average body size and gill raker number. To ensure that all populations included in the analysis had potential access to all categories of spawning habitat, populations from small and/or shallow lakes (<100 ha, <15 m maximum depth) were excluded. Altogether, 72 whitefish populations from 48 lakes filled these criteria.

Information about spawning depth and habitat were used to categorize populations as stream spawners, shallow lake spawners (depth ≤4 m) or deep lake spawners (depth >4m). The data were then analysed with multinomial regression (multinom procedure in nnet module of R^88^) using average body size and no of gill rakers as predictors and the three spawning categories as response. As the fishers’ estimates of average size could be biased by the type of gear they used, we assessed the robustness of this data by comparing individual interview data points to corresponding average sizes from our own samples. This comparison was partly based on the subset of populations that we had targeted with sampling on their spawning grounds (N=22), using non-standardized gillnet sampling (n=5), ice fishing (n=3) or hand netting (n=17, i.e. some populations were sampled with more than one method). We also included average sizes from the standardized sample fishing (not performed on spawning grounds) if the given population/ecotype could be separated from the rest of the catch by visual inspection of size- and gill raker data (n=21). Regardless of sampling method, the interview data correlated well with our sample data (Supplementary Fig. 5).

#### Genetic analyses

To identify genetic divergence indicative of reproductive isolation and to investigate the structuring of genetic diversity among and within the introduced whitefish populations, we compared neutral microsatellite genotypic data for ecotypes in 32 lakes. We performed population genetic analyses in 30 of these lakes, and extracted data from the published literature for the remaining two ^83^ (see Supplementary Table 1). 18 of the analysed lakes have whitefish populations originating from introductions between 1784 and 1985. One lake (Valsjön) has conflicting information about the introduction date, and 13 lakes have purportedly native whitefish. Individual fish were assigned to ecotype either through sampling on ecotype-specific spawning grounds or through separation of adult fish based on differences in size and morphology.

Population genetic analyses of sampled whitefish were carried out on genotypes derived from two fully overlapping marker panels comprising nine or 19 polymorphic, di- and tetranucleotide microsatellite loci. Individuals included in the 19 loci data set (36 populations, 16 lakes) formed a fully nested subset within the more extensive nine loci data set (69 populations, 30 lakes). The microsatellite loci used in this study were previously developed for the *Coregonus lavaretus* species complex^89–91^ (*ClaTet1, ClaTet3, ClaTet5, ClaTet6, ClaTet9, ClaTet1, ClaTet12, ClaTet15, ClaTet18, Cocl-Lav04, Cocl-Lav06, Cocl-Lav10, Cocl-Lav18, Cocl-Lav27, Cocl-Lav52, Cocl-Lav49, BWF2, ClaTet13, C2-157*), and were amplified in four polymerase chain reaction (PCR) multiplexes in 2.5 μl reaction volume following the PCR protocol and conditions in ^92^. PCR products were analysed using an ABI 3130XL Genetic Analyzer (Applied Biosystems Inc., Foster City, CA) and fragment lengths were analysed using GENEMAPPER^®^ 4.0 software (Applied Biosystems Inc.). Deviations from linkage equilibrium (LE) and from Hardy-Weinberg equilibrium (HWE) across ecotype samples and across loci were calculated in GENEPOP 4.5.2 ^93^ (10,000 dememorization steps, 100,000,000 Markov chain steps). *P*-values for the LE and HWE tests were corrected with the sequential Bonferroni method^94^. To reduce potential biases introduced into population genetic analyses by the presence of excessively closely related individuals (ECRs), both nine and 19 loci genotype sets were analysed in the R package RELATED^95^. Following simulations, the triadic likelihood estimator ^96^ was used to identify ECR individuals within each population. One of the individuals in each ECR pair was then excluded from all subsequent analyses.

Genetic differentiation between sympatric ecotypes was quantified as pairwise multilocus estimates of *F*_ST_, using ARLEQUIN 3.5.1.2 ^97^, with 1000 permutations to test significance. To investigate the geographic origins of within-lake genetic diversity in the introduced populations (38 ecotypes, 18 lakes), individual assignment analyses were run using STRUCTURE 2.3.4 ^98^. Parameters used: 50000 burn-in length, 500,000 MCMC chain replicates, admixture model of ancestry, correlated allele frequencies, population specific alpha prior (starting prior of 0.1). For each K, 10 independent STRUCTURE runs were carried out, up to a K of 25. For the STRUCTURE results, the true number of distinct genotypic clusters was estimated by selecting the population grouping (K) with the highest log probability of the data (ln Pr(*X*|*K*)). The STRUCTURE results were summarized and visualized using CLUMPAK^99^ and the R package POPHELPER 2.2.5 ^100^. To corroborate the STRUCTURE results, hierarchical relationships among introduced populations were reconstructed using unrooted neighbor-joining (NJ) trees of Cavalli-Sforza cord distances (*D*_*CH*_), run in PHYLIP 3.695 ^101^. Support for the recovered tree topology was estimated using 1,000 bootstrap replicates. The resulting tree was visualized in FIGTREE v1.4.2 (http://tree.bio.ed.ac.uk/software/figtree/). For both the STRUCTURE and the PHYLIP analyses, the nine loci genotype set was used for all included populations.

The results of population genetic analyses are summarized in Supplementary Table 1 (where available, only the results for 19 loci are reported). For the nine loci data set, significant locus-specific deviations from HWE were found in 36 out of 692 tests (p<0.05). Pairwise tests of linkage disequilibrium (LD) between loci were found to be significant in 79 out of 2630 tests (p<0.05). For the 19 loci data set, significant deviations from HWE were found in 45 out of 905 tests (p<0.05). LD between loci were found to be significant in 240 out of 7694 tests (p<0.05). For all LD and HWE analyses, no tests remained significant following Bonferroni correction.

For the Structure results, the population grouping with the highest log probability was found to be *K* = 16. At this K, patterns of individual cluster assignment within lakes fell into two broad categories (Supplementary Fig. 4): (i) introduction lakes without a clear signal of secondary introduction (Supplementary Fig. 4b), and (ii) introduction lakes showing signals of secondary contact between distinct genotypic clusters (Supplementary Fig. 4c).

NJ tree-based relationships among ecotypes within lakes were consistent with the patterns of individual genetic cluster assignment (Supplementary Fig. 4a). For the primary divergence lakes, most co-existing species pairs were strongly supported sister species with relatively short branch lengths (bootstrap support 100 %). Exceptions were the clades formed by Oxvattensjön/Rissjön, and Hetögeln/Murusjöen, respectively, which showed strong support (100 %) for monophyly. For Oxvattensjön/Rissjön, this reflects the introduction from the same source population ^102^, and for Hetögeln/Murusjöen, it likely reflects that Hetögeln’s dwarf, which spawns in the connecting stream, has spread upstream to Murusjöen. For the lakes included in the secondary contact category, co-existing species generally grouped closest to allopatric populations in other lakes. Only Rosången (50.2 %) and Hökvattnet (77.6 % and 63.1 %) whitefish formed monophyletic groupings, perhaps indicative of more complex secondary contact scenarios with introgression.

The primary divergence lakes were included in the chronosequence of introduced populations used in Figs. 3 and 4. Note that the dwarf/giant ecotype pair in lake Bölessjön was included in the chronosequence even though there is a third, genetically distinct ecotype that was introduced more than hundred years later than the first introduction. The inclusion of this lake was motivated by the apparent lack of introgression between the third ecotype and the original dwarf/giant ecotype pair (Supplementary Fig. 4b, probably explained by spawning segregation in space (stream vs lake) and in the timing of spawning).

#### The adaptive dynamics model

We investigated the conditions for divergence in whitefish with an adaptive dynamics approach, using a physiologically structured population model (PSPM, see refs^103–106^) in which the population has a continuous size structure and individuals reproduce continuously. Our model contains two habitats – littoral and pelagic – to which whitefish have access at all times. Each habitat has one unique resource type: macroinvertebrates are found in the littoral habitat and zooplankton in the pelagic habitat. An important difference between these resources lies in the way that resource-use efficiencies for whitefish depend on whitefish size. While the feeding efficiency for zooplankton has a hump-shaped relationship to the size of the consumer, it increases almost linearly with whitefish body size for benthic invertebrates (Supplementary Fig. 6a)^57^. Hence, large whitefish generally depend on benthic invertebrates to sustain positive growth.

However, the benefits of shifting to the benthic resource also depend on size- and habitat-specific mortality rates. Both habitats have equal background mortality rates that are unrelated to size. Because pike is a mainly littoral predator^53, 107^, pike predation is modelled as an extra, negatively size-dependent mortality rate that affects individuals feeding on the benthic resource (Supplementary Fig. 6b). Individuals allocate their time in each habitat in order to minimize the ratio between mortality rate and prey encounter rate (Supplementary Figs. 6c and d). The intake rate of a given foraging strategy is determined by resource type, resource density and individual size. In order to keep the model structure conservative and simple, there is no genetic or ecological variation among new recruits. Hence, the only way to become different from other individuals of the same size is by acquired changes in the evolving trait, namely maturation size. The model is deterministic and does not include a genetic mechanism. Thus it produces evolutionary divergence under the implicit assumption that assortative mating is present when the population reaches a branching point (or alternatively that reproduction is clonal). A detailed description of the model and parameter values is given in Supplementary Methods, Supplementary Figs. 6 and 7 and Supplementary Table 3-5.

## Acknowledgements

We thank André de Roos, Folmer Bokma and Arne Nolte for valuable comments on an earlier version of this manuscript, and the large number of local fishers and fishery managers who reported their observations in interviews and assisted in various ways during field work. We thank Emma Andersson, Jens Andersson, Sophie Bodenstein, Alex Dew, Emilija Emma, Mats-Jerry Eriksson, Johan Fahlman, Björn Fallgren, Olof Filipsson, Erik Forsberg, Tanja L. Hanebrekke, Mikael Johansson, Sandra Kero, Johan Leander, Johan Lidman, Emil Lindberg, Sofi Lundbäck, Oscar Lövbom, Martina Magnusson, Sven Norman, Johan Nyberg, Fredrik Olajos, Carmen Secco Perez, Anders Pålsson, Jan Roos and Benjamin Weigel for gillnet sampling and lab work, and Gavin Horsburgh, Lotta Ström and Rachel Tucker for technical support. This research was funded by grants from FORMAS (#2007-1149) to GE, the Swedish Research Council (#2013-5110) to GE, Biodiversa (#2012-1826) to GE and CLH, and from Göran Gustavssons Stiftelse för natur och miljö i Lappland to GÖ and GE.

## Author contributions

The study was conceived by GÖ, GE and SOÖ, and planned and coordinated by GÖ and GE. MB developed the eco-evolutionary model with ecological input from GÖ, GE and KAN and ecological/mathematical input from ÅB. MB performed the numerical simulations and the model analysis and wrote the model description with the assistance of KAN and ÅB. KBM, AGH, and KP performed genetic analyses. GÖ and SOÖ conducted interviews and collected information from different archives. GÖ, GE, SOÖ, and MP organized and undertook sample fishing surveys with help from AGH, PB and PJ, and MP and PJ performed analyses of fish in the lab as a part of their candidate- and/or master theses. GÖ, GE and PB analysed non-genetic data, and GÖ wrote the text. All authors read and commented on the manuscript before submission.

## Materials & Correspondence

Correspondence and material requests should be addressed to Gunnar Öhlund.

## Data availability

The data that support the findings of this study are available from the corresponding author upon reasonable request.

