## Supplementary Information for "Ecological speciation in European whitefish is driven by a large-gaped predator"

**Supplementary Methods: An adaptive dynamics model of whitefish divergence**

### 1.1 Model overview

For the theoretical analysis, we use a physiologically structured population model (PSPM)[1, 2], which is based on the models described in Claessen et al.[3] and Andersson[4]. We developed the population model to account for predation- and resource-dependent habitat choice and evolution of size at maturation. Individuals are characterized by body size and reside in two separate habitats, corresponding to the pelagic and the littoral zones of a lake. The pelagic habitat has a zooplankton resource, while the littoral habitat has a macroinvertebrate resource and an elevated risk of predation. Individuals balance resource intake rate and mortality risk when choosing the habitat in which to forage. In our model, a niche shift arises as a consequence of adaptive foraging (i.e. individuals shift from feeding on zooplankton to invertebrates as they grow larger). Individuals initially allocate all their energy to maintenance and growth, but after maturation most of the energy is used for maintenance and reproduction. Predation in the littoral habitat is incorporated as a density-independent mortality that decreases with size and approaches zero at the predator's maximum gape size.

### 1.2 Parameters

Most of the model parameters (Supplementary Table 3) are estimated for Arctic charr [4] rather than whitefish. We believe that this is justified, as the two species are relatively similar in terms of basic vital rates. One important difference is that whitefish is a more efficient planktivore than charr[5], and continues to feed on zooplankton up to a larger body size than is observed for charr[6, 7]. Thus, we used 15 g as the optimal body weight for feeding on zooplankton, compared to the value of 7.15 g that has been used for charr[8]. The parameter $q$, which represents the survival of the eggs and larvae, has not been measured empirically. We set the survival rate to a relatively low value since egg cannibalism and other types of juvenile mortality are reported to be high in whitefish populations[9]. It is noteworthy that changing the survival parameter $q$ has the same effects as changing the energy cost of producing one egg (now set to 1). The constants controlling energy allocation between reproduction, growth and metabolism were selected to produce a marked change in energy allocation when individuals mature. The metabolic scaling exponent $\alpha$ was chosen as the median for teleost fish[10] .

As the model is parameter-rich we did not perform an exhaustive examination of the parameter space. However, as an indication of the robustness of the main result, i.e., that predation results in evolutionary divergence, we provide approximate ranges of parameter values within which this conclusion holds (Supplementary Table 5). The examination of robustness was limited to parameters that are poorly known.

### 1.3 Ecological dynamics

Brief descriptions of the model parameters, variables, and functions are given in Supplementary Tables 3-4 and Supplementary Fig. 6. The change in the prey density $n(s,t)$ of individuals of size $s > s_0$ at time $t$ is given by the McKendrick-von Foerster equation,

$$\frac{\partial}{\partial t}n(s,t) + \frac{\partial}{\partial s}g(s,\mathbf{R})n(s,t) = -d(s,\mathbf{R})n(s,t)$$

with the inflow of newborn individuals given by the boundary condition

$$g(s_0,\mathbf{R})n(s_0,t) = \int_{s_m}^{\infty} b(s,\mathbf{R})n(s,t)ds$$

in which $s_0$ is the size at birth and $s_m$ is the maturation size. We denote the zooplankton density in the pelagic habitat by $R_1$ and the macroinvertebrate density in the littoral habitat by $R_2$. The two resources follow semi-chemostat dynamics, i.e., the dynamics of the resource $R_i$ is given by

$$\frac{d\mathbf{R}_i}{dt} = r_i(K_i - \mathbf{R}_i) - V_i^{-1}\int_{s_0}^{\infty} f_i\,(s,\mathbf{R})n(s,t)ds$$

where $\mathbf{R}$=$(R_1, R_2)$ denotes the resource vector. Numerical simulations of population dynamics were carried out using the Escalator Boxcar Train method[1, 11, 12].

### 1.4 Adaptive foraging

The ability of fishes to estimate the mortality risk from predators and choose their habitat accordingly has been documented empirically in several species[13-15]. We assume that individuals account for resource availability and mortality risk when selecting a habitat. Specifically, for each size, individuals strive to minimize the ratio of mortality rate to resource encounter rate. We chose this criterion because it is consistent with empirical evidence showing that fishes can balance resource abundance and predation risk[16]. There is no cost of moving between the two habitats.

We describe adaptive foraging by the fraction of time, $T_i(s)$, an individual spends foraging in habitat $i$, which we assume is given by

$$T_i(s,\mathbf{R}) = \frac{(a_i(s)\mathbf{R}_i/\mu_i(s))^\theta}{(a_1(s)\mathbf{R}_1/\mu_1(s))^\theta + (a_2(s)\mathbf{R}_2/\mu_2(s))^\theta}$$

The attack rates on the resources and the mortality rate in each habitat are illustrated in Supplementary Figs. 6a and b. The parameter $\theta$ describes the strength of habitat selection, where $\theta = 0$ indicates random habitat choice, $\theta = 1$ means that individuals chose habitat proportionally to the mortality/resource encounter ratio, and $\theta > 1$ indicates that individuals favour the habitat with the higher ratio. Supplementary Figs. 6c and d shows the resulting size-dependent habitat use when the habitat selection rule is applied. As the fish grow to a large size, their inability to feed on zooplankton forces a switch to the littoral habitat and a diet of macroinvertebrates.

### 1.5 Energy budget, growth and reproduction

We adopt a net production energy-budget model where metabolic requirements are met first, and the remaining energy is allocated to growth and reproduction. The resource intake rate is given by a Holling Type-II functional response. Assuming the same conversion efficiency, $\varepsilon$, in the two habitats, the total energy intake rate from both resources is $\varepsilon f(s, \mathbf{R})$, cf. Supplementary Tables 3 and 4. We assume that the metabolic requirements scale allometrically with body weight, $W$. The energy remaining after the metabolic needs have been covered equals

$$\varepsilon f(s, \mathbf{R}) - \beta_1 W^{\beta_2}$$

This energy is divided between growth and reproduction, where the fraction of energy allocated to growth is described by the decreasing function, $K(s) = K_{s_m}(s)$, which declines most steeply at the size when maturation is reached. Should the energy required for maintenance exceed the total energy available, we assume that the individual dies from starvation.

### 1.6 Evolutionary dynamics

Adaptive dynamics techniques[17, 18] were used to investigate the dynamics of the system. Specifically, we assume that the mean population trait value $s_m$ evolves according to a gradient dynamical system known in the adaptive-dynamics literature as the canonical equation[12]:

$$\frac{ds_m}{dt} = \frac{1}{2}\rho\sigma^2 N(s_m) D(s_m)$$

Here, $D$ is the selection gradient, $\rho$ is the mutation probability, $s^2$ is the variance of the mutation distribution, and $N(s_m)$ is the equilibrium population size. Since $\rho$ and $\sigma^2$ are positive, the factor $\rho\sigma^2/2$ only scales the rate of evolutionary change and, therefore, does not affect the location of the evolutionarily stable maturation size. The selection gradient is given by

$$D(s_m) = \left.\frac{\partial I_{s_m}}{\partial s'_m}\right|_{s'_m = s_m}$$

where $I_{s_m}(s'_m)$ denotes the invasion fitness[19]. The invasion fitness is the long-term exponential per capita growth rate of an initially rare mutant with trait $s'_m$ in a monomorphic population in which the residents mature at size $s_m$.

At demographic equilibrium, the resource levels, $\mathbf{R}$, set by the resident population determine the environment experienced by a mutant. Let $\Pr(s)$ denote the probability that a newborn mutant maturing at size $s'_m$ survives until it reaches size $s$, in a population consisting of individuals maturing at size $s_m$, which is given by

$$\Pr(s) = \exp\left( - \int_{s_0}^{s} \frac{d(l, \mathbf{R})}{g_M(l, \mathbf{R})} dl \right)$$

where $g_M(l, \mathbf{R})$ is the ontogenetic growth rate of the mutant and $d(l, \mathbf{R})$ is the death rate determined by the resident population. We can then calculate the basic reproduction ratio, $R_0(s_m, s'_m)$, for the mutant as

$$R_0(s_m, s'_m) = \int_{s'_m}^{\infty} \frac{b_M(s, \mathbf{R})}{g_M(s, \mathbf{R})} \Pr(s)\, ds$$

where $b_M(s, \mathbf{R})$ is the birth rate of the mutant. Note that $g_M(s, \mathbf{R})$ and $b_M(s, \mathbf{R})$, only differ from the resident's growth and birth rate through a change of maturation size in $\kappa(s)$, cf. Supplementary Table 4. The invasion fitness is, in general, given by the dominant Lyapunov exponent of the mutant's dynamics, but since this is difficult to calculate in practice we use the sign equivalent version, $I_{s_m}(s'_m) = ln[R_0(s_m, s'_m)]$. Then the selection gradient equals

$$D(s_m) = \left. \frac{\partial \ln[R_0(s_m, s'_m)]}{\partial s'_m} \right|_{s'_m = s_m} = \left. \frac{\partial R_0(s_m, s'_m)}{\partial s'_m} \right|_{s'_m = s_m}$$

since $R_0(s_m, s_m) = 1$ at demographic equilibrium. A strategy $s_m$ is called a evolutionarily singular strategy if $D(s_m) = 0$. If the invasion fitness has a maximum at a convergence stable singular strategy $s_m$ it is an evolutionarily stable strategy, and if the invasion fitness has a minimum at $s_m$, it is an evolutionary branching point.

### 1.7 Simulation

We simulated the system as the evolution of size at maturation in a monomorphic population. Before simulation is initiated, there can be one or two convergence stable singular strategies depending on predation intensity (as illustrated by the pairwise invasibility plots in Supplementary Fig. 7). When there are two convergence stable singular strategies, the larger one is evolutionarily stable, and the smaller one is either a branching point or evolutionarily stable, depending on predation intensity (Supplementary Figs. 6b and c). The simulation was initiated with a monomorphic population having a maturation size close to the smallest singular strategy, which then was allowed to evolve until it reached a singular strategy. If the singular strategy was a branching point, the monomorphic population was replaced by two populations; one maturing at a slightly larger size and one at a slightly smaller size than the monomorphic population that they replaced. They were then allowed to evolve together until evolutionary stability was reached. Fig. 6a in the main text shows the evolutionary outcome.

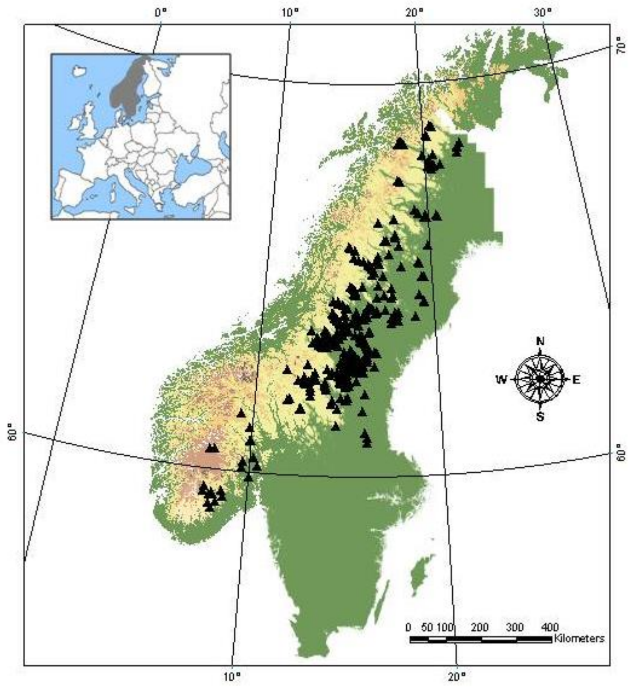


**Supplementary Figure 1. Map of Scandinavia showing the geographical distribution of the 357 lakes in our data set.**

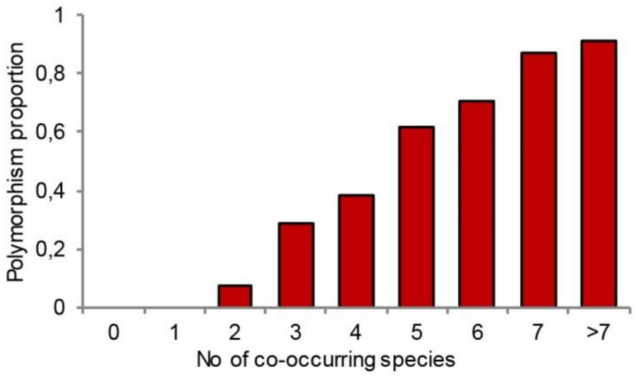


**Supplementary Figure 2. Dwarf and giant ecotypes of whitefish develop in relatively species rich lakes.** The likelihood of finding two or more ecotypes of whitefish increases with fish species richness (logistic regression: coefficient±SE=0.80 ±0.09, Z=8.4, N=350, $p<10^{-16}$). The relationship is also positive if richness is standardized with respect to lake area (coefficient±SE=0.70±0.19, Z=3.77, p=0.0002). Richness was standardized by dividing with $Area^c$, where the exponent (c=0.166) was determined by regressing log(species richness) on log(area).

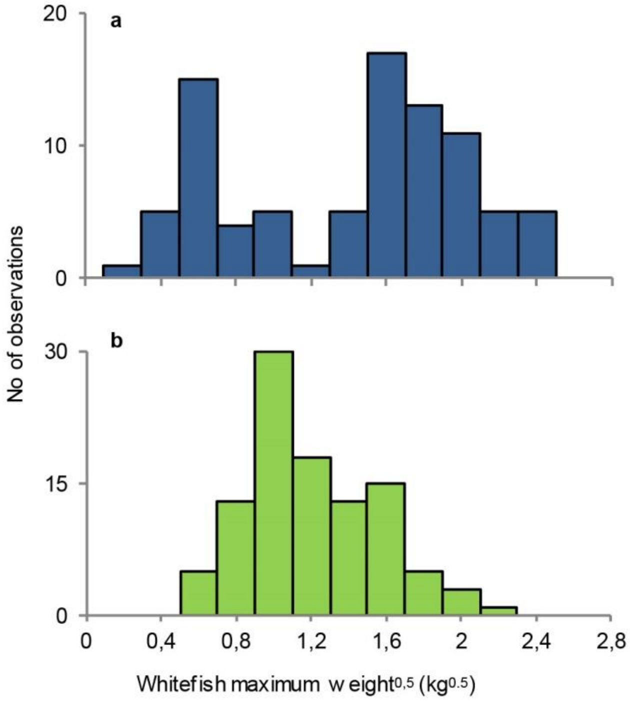


**Supplementary Figure 3. Pike presence induces either dwarfs or giants in monomorphic populations.** Frequency distribution of maximum weights of monomorphic whitefish populations in a) lakes with pike (blue colour, n=87) and b) in lakes without pike (green colour, n=103). Cluster analyses provide support for two clusters in pike lakes (ΔBIC=25.3), and a single cluster in lakes without pike (ΔBIC=4.3).

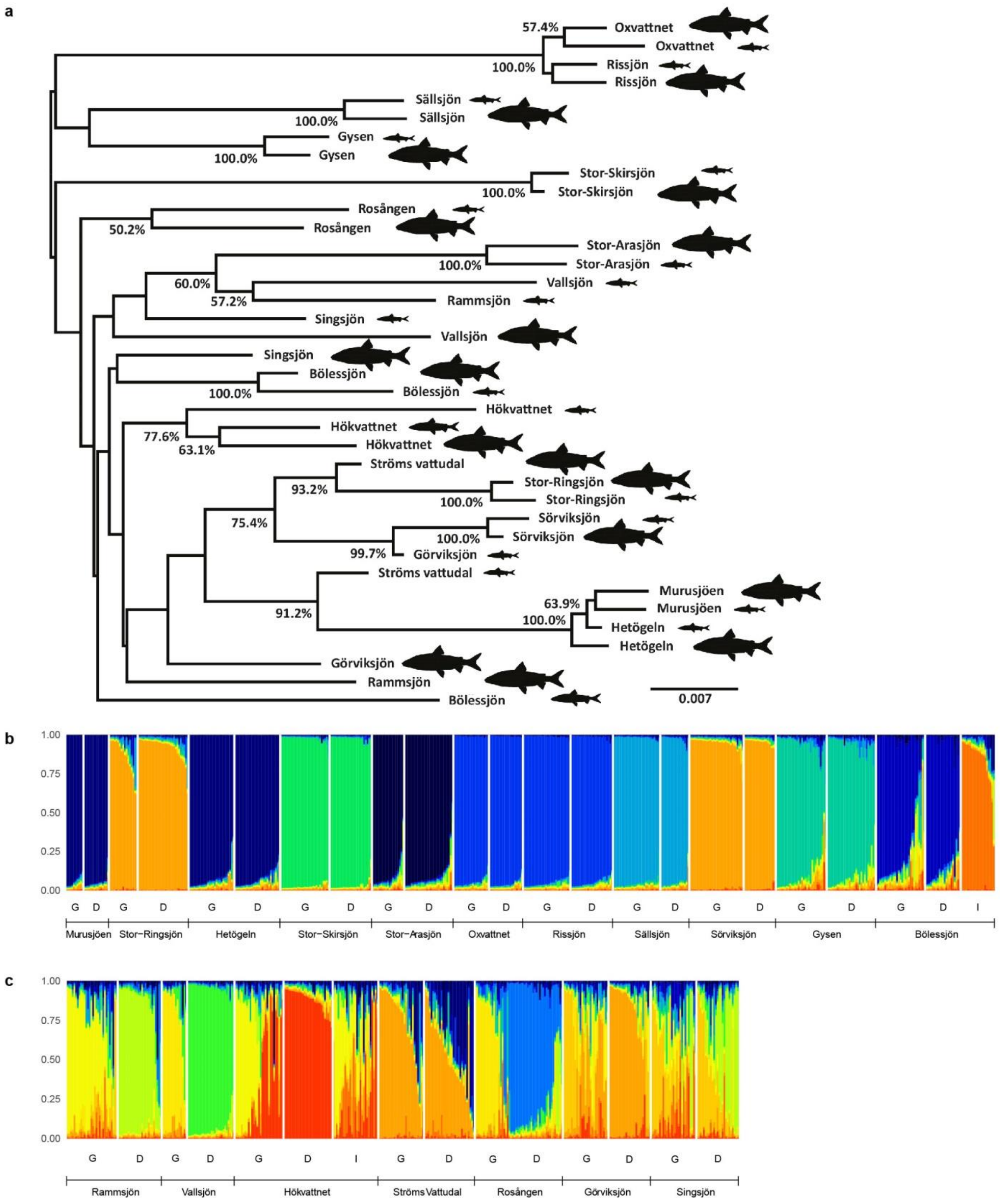


**Supplementary Figure 4. Selection of introduced populations for the chronosequence.** Genetic data that motivated our decision to include or exclude populations from the chronosequence of lakes with pike and introduced whitefish that were used in Figures 3 and 4 (pike present category). a) Population-based unrooted neighbor-joining tree of Cavalli-Sforza chord distances ($D_{CH}$) showing the genetic relationships among whitefish from the different lakes. Numerical values refer to bootstrap support > 50% (1000 iterations). Images of different size at the tips indicate the assigned ecotype: Giant, Dwarf or Intermediate. b) and c) are structure plots showing the populations that were (b) included in, and (c) excluded from the chronosequence. G=giant-, D=dwarf-, and I=intermediately sized ecotypes.

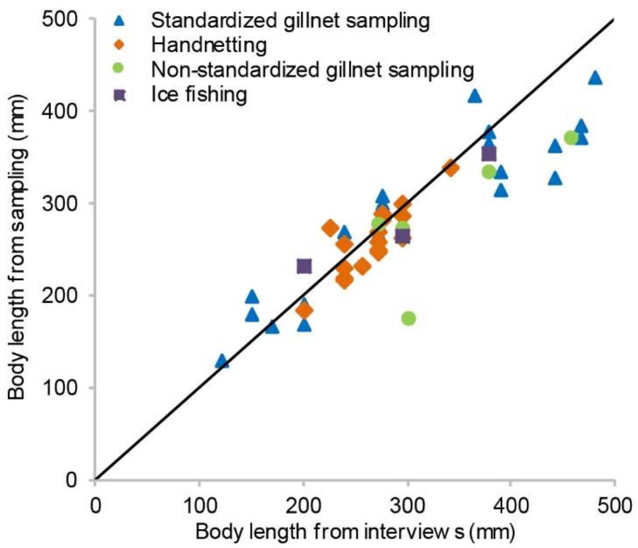


**Supplementary Figure 5. Validation of interview data on mean body length.** Average body length from our sampling of whitefish populations are compared with corresponding estimates reported by local fishers. The sampling methods include standardized gillnet sampling outside of the spawning season (N=21), as well as hand netting (N=17), gill netting (N=5) and ice fishing (N=3) on spawning grounds.

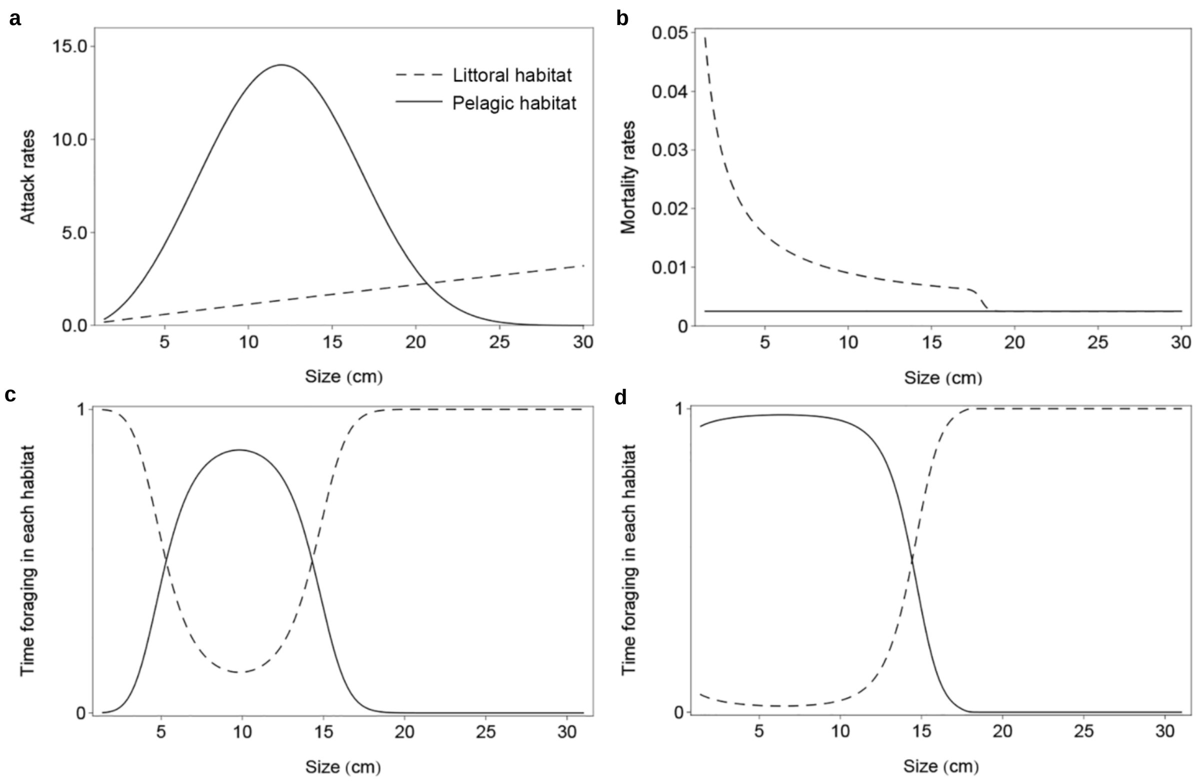


**Supplementary Figure 6. Size-dependent ecological rates in the different habitats and effects of predation risk on habitat choice in the model.** a) Attack rate on pelagic and littoral resources. b) Mortality rates in the pelagic and littoral habitats. c) and d) Fraction of time spent foraging in the pelagic and littoral habitats without predation (c) and with predation (d). In this simulation, maturation size is 14 cm, maximum prey size eaten by the predator is 18 cm, and the predation intensity is 80%. The functions are specified in Supplementary Table 4.

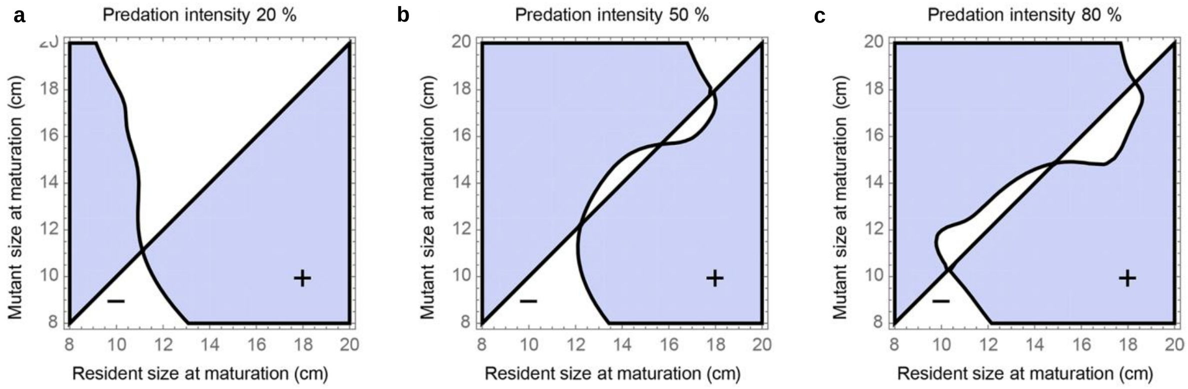


**Supplementary Figure 7. Pairwise invasibility plots for three different predation intensities.** a) Low predation intensity yields one evolutionarily stable maturation size. b) Medium predation intensity yields two convergence stable singular strategies, where the larger one is evolutionarily stable and the smaller one is a branching point. c) High predation intensity yields two evolutionarily stable maturation sizes.

**Supplementary Table 1.** Genetic data and geographic location for the lakes (n=32) where we present population genetic analyses. For lakes with more than two ecotypes, we present the genetic distance between the largest and the smallest ecotype. Two of the lakes, Stor-Ringsjön and Murusjøen, have whitefish populations that are too young to be included in the interview-based analysis underlying Figure 1b, and hence, also to be used in the validation of said interview-based results that is presented in the results section.

| Lake | Ecotype | Latitude | Longitude | Year of introduction | # μ-sat markers | # Pop Gen | $F_{ST}$ | HWE (*P*-Value) |
|---|---|---|---|---|---|---|---|---|
| Bodsjön | Giant | 62.8527 | 15.4145 | | 19 | 22 | 0.054* | 0.97 |
| | Intermediate | | | | | 26 | | 0.50 |
| | Dwarf | | | | | 26 | | 0.83 |
| Bölessjön[†] | Giant | 62.9263 | 14.8418 | 1825 | 9 | 29 | 0 | 0.21 |
| | Intermediate | | | | | 20 | | 0.94 |
| | Dwarf | | | | | 21 | | 0.42 |
| Femunden[‡] | Bay spawner[‡] | 61.9349 | 11.8633 | | 6 | 45 | 0.120* | <0.05 |
| | Deep spawner[‡] | | | | | 182 | | <0.05 |
| | Stream spawner[‡] | | | | | 146 | | <0.05 |
| | Shallow spawner[‡] | | | | | 91 | | <0.05 |
| Gysen[†] | Giant | 63.6414 | 14.3934 | 1830 | 19 | 30 | 0.007* | 0.88 |
| | Dwarf | | | | | 28 | | 0.23 |
| Gåxsjön | Giant | 63.6729 | 15.1012 | | 19 | 14 | 0.057* | 0.24 |
| | Dwarf | | | | | 20 | | 0.91 |
| Görvikssjön | Giant | 63.5989 | 15.7038 | 1845 | 9 | 27 | 0.025* | 0.12 |
| | Dwarf | | | | | 25 | | 0.76 |
| Hetögeln[†] | Giant | 64.3884 | 14.4208 | 1960 | 9 | 27 | 0 | 0.73 |
| | Dwarf | | | | | 27 | | 0.88 |
| Hotagen | Giant | 63.7890 | 14.6680 | | 19 | 19 | 0.068* | 0.95 |
| | Dwarf | | | | | 30 | | 0.93 |
| Hökvattnet | Giant | 63.8834 | 14.8386 | 1865 | 9 | 29 | 0.064* | 0.04* |
| | Intermediate | | | | | 27 | | 0.14 |
| | Dwarf | | | | | 29 | | 0.04* |
| Idsjön | Giant | 62.8180 | 15.7121 | | 19 | 26 | 0.055* | 0.90 |
| | Intermediate | | | | | 29 | | 0.47 |
| | Dwarf | | | | | 22 | | 0.58 |
| Ismunden | Giant | 63.1474 | 15.1963 | | 19 | 18 | 0.102* | 0.64 |
| | Dwarf | | | | | 20 | | 0.95 |
| Isteren[‡] | ”Normal” | 61.9096 | 11.7774 | | 6 | 30 | 0.085* | >0.05 |
| | ”Dwarf” | | | | | 30 | | >0.05 |
| Locknesjön | Giant | 62.9198 | 14.9395 | | 19 | 29 | 0.022* | 0.45 |
| | Intermediate | | | | | 15 | | 0.93 |
| | Dwarf | | | | | 25 | | 0.63 |
| Långvattnet | Giant | 62.9629 | 15.1757 | | 19 | 7 | 0.051* | 1.00 |
| | Dwarf | | | | | 23 | | 0.05* |
| Murusjøen | Giant | 64.4597 | 14.1022 | 1990 | 19 | 10 | 0.011 | 0.95 |
| | Dwarf | | | | | 15 | | 0.54 |
| Oxvattensjön[†] | Giant | 64.0905 | 17.8637 | 1926 | 9 | 21 | 0.001 | 0.54 |
| | Dwarf | | | | | 20 | | 0.44 |
| Rammsjön | Giant | 62.3789 | 14.5869 | 1930 | 9 | 30 | 0.064* | 0.21 |
| | Dwarf | | | | | 26 | | 0.79 |
| Revsundssjön | Giant | 62.8140 | 15.3534 | | 19 | 29 | 0.055* | 0.91 |
| | Intermediate | | | | | 23 | | 1.00 |
| | Dwarf | | | | | 27 | | 0.69 |
| Rissjön[†] | Giant | 64.0289 | 17.8095 | 1926 | 9 | 28 | 0 | 0.88 |
| | Dwarf | | | | | 25 | | 0.45 |
| Rosången | Giant | 62.2980 | 14.9040 | 1860 | 9 | 21 | 0.055* | 0.99 |
| | Dwarf | | | | | 31 | | 0.60 |
| Singsjön | Giant | 63.1206 | 15.0922 | 1784 | 9 | 27 | 0.036* | 0.67 |
| | Dwarf | | | | | 25 | | 0.31 |
| Skåsjön | Giant | 62.8035 | 15.7156 | | 19 | 7 | 0.041* | 1.00 |
| | Intermediate | | | | | 22 | | 0.96 |
| | Dwarf | | | | | 29 | | 0.65 |
| Stor-Arasjön[†] | Giant | 64.5986 | 17.6004 | 1937 | 9 | 19 | 0.002 | 0.35 |
| | Dwarf | | | | | 29 | | 0.30 |
| Stor-Ringsjön[†] | Giant | 64.2646 | 14.9974 | 1984 | 19 | 16 | 0 | 0.63 |
| | Dwarf | | | | | 30 | | 0.96 |
| Stor-Skirsjön[†] | Giant | 64.0659 | 16.0843 | 1945 | 9 | 29 | 0.010 | 0.77 |

| | | | | | | | | |
|---|---|---|---|---|---|---|---|---|
| | Dwarf | | | | | 25 | | 0.42 |
| Storvindeln | Giant | 65.6369 | 17.4507 | | 19 | 23 | 0.036* | 0.17 |
| | Interm. dens. rakered | | | | | 28 | | 0.80 |
| | Interm. spars. rakered | | | | | 25 | | 0.85 |
| | Dwarf | | | | | 21 | | 0.38 |
| Ströms Vattudal | Giant | 63.9303 | 15.6104 | 1854 | 9 | 27 | 0.015* | 0.06 |
| | Dwarf | | | | | 30 | | 0.56 |
| Sundsjön | Giant | 62.9629 | 15.1757 | | 9 | 28 | 0.025* | 0.90 |
| | Dwarf | | | | | 8 | | 0.48 |
| Sällsjön† | Giant | 63.2376 | 13.6940 | 1870 | 19 | 28 | 0.024* | 0.05 |
| | Dwarf | | | | | 9 | | 0.97 |
| Sörvikssjön† | Giant | 63.5520 | 15.8496 | 1845 | 19 | 32 | 0.001 | 0.60 |
| | Dwarf | | | | | 19 | | 0.83 |
| Valsjön | Giant | 64.0198 | 14.2074 | <1900§ | 19 | 28 | 0.119* | 0.99 |
| | Dwarf | | | | | 28 | | 0.02* |
| Vallsjön | Giant | 62.9226 | 16.0656 | 1915 | 9 | 15 | 0.073* | 0.97 |
| | Dwarf | | | | | 28 | | 0.83 |

* $P < 0.05$

† Introduced populations of whitefish that were included in the chronosequence of lakes shown in Figs. 3 and 4 (pike present category).

‡ Data from Østbye et al. 2005 [20].

§ Conflicting information about introduction date

**Supplementary Table 2.** Differences between coexisting dwarf and giant whitefish ecotypes in the number of gill rakers (n=72). Two of the lakes, Stor-Ringsjön and Murusjøen, have whitefish populations that are too young to be included in the interview-based analysis underlying Figure 1b, and hence, also to be used in the validation of said interview-based results that is presented in the results section.

| Lake | Year of introduction | Latitude | Longitude | Gill raker difference | t-value | p-value | No of Obs. | Reference |
|---|---|---|---|---|---|---|---|---|
| Alsensjön | | 63.3241 | 14.1493 | 8.5 | 9.8 | <0.0001 | 67 | [21] |
| Ansjön | | 62.9764 | 16.0890 | 2.3 | 18.5 | <0.0001 | 3192 | [22] |
| Bodsjön | | 62.8662 | 14.9831 | 10.1 | 13.2 | <0.0001 | 69 | [22] |
| Bodsjön | | 62.8527 | 15.4145 | 8.6 | 19.3 | <0.0001 | 60 | this study |
| Bomsjön | 1895 | 64.6009 | 17.1052 | 0.72 | 1.58 | 0.12 | 121 | this study |
| Bölessjön† | 1825 | 62.2372 | 17.3906 | 2.33 | 4.57 | <0.0001 | 122 | this study |
| Dikasjön | | 65.2145 | 16.0419 | 13.6 | 29.4 | <0.0001 | 158 | [23] |
| Femund | | 61.9178 | 11.9445 | 16.6 | 63.1 | <0.0001 | 219 | [22] |
| Flåsjön | | 64.1308 | 15.9169 | 6.6 | 10.0 | <0.0001 | 43 | this study |
| Gesunden | | 63.1460 | 16.0698 | 4.6 | 6.8 | <0.0001 | 105 | [21] |
| Gysen† | 1830 | 63.6414 | 14.3934 | 1.53 | 5.48 | <0.0001 | 244 | this study |
| Gåxsjön | | 63.6729 | 15.1012 | 19.2 | 39.1 | <0.0001 | 71 | this study |
| Gärdessjön | 1908 | 63.5596 | 13.7502 | 0.30 | 0.59 | 0.24 | 65 | this study |
| Görvikssjön | 1845 | 63.5989 | 15.7038 | 7.0 | 9.5 | <0.0001 | 79 | this study |
| Hammerdalssjön | | 63.5384 | 15.4453 | 16.2 | 28.6 | <0.0001 | 40 | [21] |
| Helgesjön | 1865 | 63.3400 | 13.3125 | 14.2 | 19.4 | <0.0001 | 98 | [21] |
| Hetögeln† | 1960 | 64.3884 | 14.4208 | 0.28 | 0.64 | 0.28 | 117 | this study |
| Hornavan | | 66.0593 | 17.8727 | 13.9 | 40.0 | <0.0001 | 425 | [22] |
| Hotagen | | 63.7890 | 14.6680 | 3.9 | 8.0 | <0.0001 | 140 | [23] |
| Härjåsjön | | 61.9160 | 14.1969 | 15.1 | 37.1 | <0.0001 | 90 | [22] |
| Hökvattnet | 1865 | 63.8834 | 14.8386 | 1.6 | 2.25 | 0.012 | 76 | this study |
| Idsjön | | 62.8180 | 15.7121 | 5.6 | 13.4 | <0.0001 | 102 | this study |
| Ismunden | | 63.1474 | 15.1963 | 19.6 | 39.6 | <0.0001 | 93 | this study |
| Laisan | | 65.9918 | 17.1964 | 24.6 | 43.1 | <0.0001 | 180 | [22] |
| Landösjön | | 63.5477 | 14.2856 | 4.4 | 39.9 | <0.0001 | 1862 | [21] |
| Locknesjön | | 62.9198 | 14.9395 | 18.5 | 37.5 | <0.0001 | 168 | this study |
| Lossen | | 62.4038 | 12.9249 | 3.5 | 20.2 | <0.0001 | 574 | [22] |
| Långvattnet | | 65.1802 | 16.5435 | 4.4 | 6.2 | <0.0001 | 58 | this study |
| Lännässjön | | 62.6315 | 14.1325 | 10.9 | 6.5 | <0.0001 | 17 | this study |
| Murusjøen | 1975 | 64.4597 | 14.1022 | 0.17 | 0.24 | 0.81 | 53 | this study |
| Näkten | | 62.9118 | 14.5694 | 17.7 | 82.4 | <0.0001 | 303 | [21] |
| Näldsjön | | 63.3460 | 14.2419 | 7.1 | 14.9 | <0.0001 | 276 | [21] |
| Ockesjön | | 63.2782 | 13.8612 | 2.5 | 7.5 | <0.0001 | 174 | [21] |
| Ormsjön | | 64.3393 | 16.1124 | 3.6 | 11.8 | <0.0001 | 348 | [22] |
| Orrmosjön | | 61.8374 | 14.1175 | 16.2 | 32.5 | <0.0001 | 85 | [22] |
| Orten-Randsjön | | 62.2565 | 13.8307 | 9.9 | 12.1 | <0.0001 | 55 | [22] |
| Ottsjön | | 63.7073 | 14.9500 | 17.4 | 26.0 | <0.0001 | 234 | [21] |
| Oxvattensjön† | 1926 | 64.0905 | 17.8637 | 0.25 | 0.46 | 0.64 | 119 | this study |
| Parkijaure | | 66.7448 | 19.2048 | 22.8 | 63.3 | <0.0001 | 287 | [22] |
| Rammsjön | 1930 | 62.3789 | 14.5869 | 7.5 | 13.1 | <0.0001 | 68 | this study |
| Randijaure | | 66.6579 | 19.4000 | 14.7 | 22.1 | <0.0001 | 81 | [24] |
| Revsundssjön | | 62.8140 | 15.3534 | 6.8 | 18.8 | <0.0001 | 82 | this study |
| Rissjön† | 1926 | 64.0289 | 17.8095 | 0.81 | 1.67 | 0.10 | 76 | this study |
| Rosången | 1860 | 62.2980 | 14.9040 | 14.1 | 19.0 | <0.0001 | 77 | this study |
| Sicksjön | | 63.0326 | 63.0302 | 11.4 | 50.5 | <0.0001 | 907 | [22] |
| Singsjön | 1784 | 63.1206 | 15.0922 | 10.0 | 18.7 | <0.0001 | 129 | this study |
| Skikkisjaure | | 65.1192 | 16.4243 | 17.6 | 36.9 | <0.0001 | 113 | [25] |
| Skåsjön | | 62.8035 | 15.7156 | 14.1 | 15.3 | <0.0001 | 81 | this study |
| Stor-Arasjön† | 1937 | 64.5986 | 17.6004 | 0.68 | 1.12 | 0.27 | 63 | this study |
| Stora Skeppsträsk | | 65.1857 | 18.9198 | 10.3 | 24.1 | <0.0001 | 443 | [25] |
| Storavan | | 65.6740 | 18.1433 | 12.5 | 51.8 | <0.0001 | 259 | [22] |

| | | | | | | | | |
|---|---|---|---|---|---|---|---|---|
| Stor-Juktan | | 65.3210 | 17.3313 | 8.5 | 19.3 | <0.0001 | 77 | [26] |
| Storsjön | | 63.2983 | 14.4601 | 15.7 | 46.5 | <0.0001 | 169 | [27] |
| Stor-Skirsjön† | 1945 | 64.0659 | 16.0844 | 0.25 | 0.66 | 0.51 | 126 | this study |
| Stor-Ringsjön† | 1985 | 64.2646 | 14.9974 | 0.61 | 1.13 | 0.27 | 54 | this study |
| Storuman | | 65.0946 | 17.1007 | 21.2 | 114.2 | <0.0001 | 958 | [22] |
| Storvindeln | | 65.6369 | 17.4507 | 16.3 | 38.9 | <0.0001 | 119 | this study |
| Ströms Vattudal | 1860 | 63.9303 | 15.6104 | 2.4 | 4.2 | 0.0009 | 113 | this study |
| Sundsjön | | 62.9629 | 15.1757 | 15.6 | 17.1 | <0.0001 | 41 | this study |
| Sällsjön† | 1870 | 63.2376 | 13.6939 | 2.71 | 5.95 | <0.0001 | 142 | this study |
| Sörvikssjön† | 1845 | 63.5520 | 15.8496 | 1.47 | 3.23 | 0.0017 | 108 | this study |
| Tåsjön | | 64.1607 | 16.0072 | 9.1 | 7.1 | <0.0001 | 30 | this study |
| Uddjaure | | 65.8464 | 18.0429 | 16.8 | 12.4 | <0.0001 | 15 | this study |
| Vallsjön | 1915 | 62.9226 | 16.0656 | 10.9 | 15.0 | <0.0001 | 51 | this study |
| Valsjön | <1900§ | 64.0198 | 14.2074 | 0.76 | 2.34 | 0.021 | 98 | this study |
| Venjanssjön | | 60.8357 | 14.1193 | 3.7 | 8.4 | <0.0001 | 106 | [22] |
| Vikarsjön | | 62.3750 | 13.7589 | 7.9 | 15.9 | <0.0001 | 92 | [22] |
| Vojmsjön | | 64.8618 | 16.7438 | 16.3 | 130.5 | <0.0001 | 1861 | [22] |
| Volgsjön | | 64.5501 | 16.7145 | 13.9 | 28.8 | <0.0001 | 431 | [25] |
| Östra Vattnan | | 62.3339 | 12.6695 | 10.1 | 17.0 | <0.0001 | 28 | [22] |
| Över-Särvsjön | | 62.6319 | 13.1551 | 15.6 | 36.9 | <0.0001 | 177 | [22] |
| Övsjön | | 63.0302 | 15.9887 | 1.2 | 3.2 | 0.0016 | 174 | [22] |

§ Conflicting information about introduction date

† Introduced populations of whitefish that were included in the chronosequence of lakes shown in Figs. 3 and 4 (pike present category).

**Supplementary Table 3.** Description of the model parameters. Superscript numbers within parentheses refers to Supplementary References.

| Constant | Value | Description | Unit |
|---|---|---|---|
| Prey physical parameters | | | |
| $s_0$ | 1.4 [4] | Size at birth | cm |
| $l_1$ | 5.03 [4] | Length to weight relation | cm $g^{-l_2}$ |
| $l_2$ | 0.32 [4] | Length weight exponent | – |
| $x_1$ | 5.33 [4] | Handling time constant | day $g^{-(1+x_2)}$ |
| $x_2$ | -0.66 [4] | Handling time exponent | – |
| $b_1$ | 0.033 [28] | Metabolic rate constant | $day^{-1} g^{-(1+b_2)}$ |
| $b_2$ | 0.7 [12] | Metabolic rate exponent | – |
| $e$ | 0.61 [4] | Conversion coefficient | – |
| $h$ | 1 | Energy cost for producing one egg | g |
| $q$ | 0.02 | Probability of surviving the larval stage | – |
| $k$ | 5 | Energy allocation slope constant | $cm^{-1}$ |
| $k_j$ | 1 | Juvenile energy allocation for growth | – |
| $k_a$ | 0.04 | Adult energy allocation for growth | – |
| $q$ | 5 | Strength of habitat selection | – |
| Pelagic habitat (Habitat 1) | | | |
| $r_1$ | 0.1 [4] | Growth rate of the zooplankton resource $R_1$ | $day^{-1}$ |
| $K_1$ | 1.0 [29] | Carrying capacity of resource $R_1$ | g $m^{-3}$ |
| $V_1$ | $10^6$ [28] | Volume of the pelagic habitat | $m^3$ |
| $\hat{a}_1$ | 14.0 [4] | Maximum attack rate on resource $R_1$ | $m^3$ $d^{-1}$ |
| $a_1$ | 0.65 [4] | Attack rate exponent for $R_1$ | – |
| $w_0$ | 15 | Weight at optimal attack rate on resource $R_1$ | g |
| $m_1$ | 0.0025 | Background mortality rate in the pelagic habitat | $day^{-1}$ |

Littoral habitat (Habitat 2)

| | | | |
|---|---|---|---|
| $r_2$ | 0.1 [4] | Growth rate of macroinvertebrate resource $R_2$ | day$^{-1}$ |
| $K_2$ | 28.0 [30] | Carrying capacity of resource $R_2$ | g m$^{-2}$ |
| $V_2$ | 50000 [28] | Area of the littoral habitat | m$^2$ |
| $\hat{a}_2$ | 0.6 [4] | Attack rate constant for resource $R_2$ | m$^2$ day$^{-1}$ |
| $a_2$ | 0.3 [4] | Attack rate exponent $R_2$ | – |
| $m_2$ | 0.0025 | Background mortality rate in the littoral habitat | day$^{-1}$ |
| $k_p$ | 5 | Predation slope constant at maximum gape size | cm$^{-1}$ |
| $g_p$ | 1 | Predation mortality exponent | – |
| $l_p$ | Varied | Maximum predator gape size | cm |
| $p$ | Varied | Predator attack rate constant | day$^{-1}$ |

**Supplementary Table 4.** Description of the model variables and functions.

| Variable | Description | Unit |
|---|---|---|
| $s$ | Size | cm |
| $s_m$ | Maturation size | cm |
| $n(s,t)$ | Population size distribution at time $t$ | cm$^{-1}$ |
| $R_1$ | Density of the pelagic resource | g m$^{-3}$ |
| $R_2$ | Density of the littoral resource | g m$^{-2}$ |

**Function**

| | | |
|---|---|---|
| $l(w) = l_1 w^{l_2}$ (below $w = l^{-1}(s)$) | Length of an individual with weight $W$ gram | cm |
| $k(s) = k_j - \dfrac{(k_j - k_a)}{1+\exp[-k(s-s_m)]}$ | Fraction of energy channelled to growth | – |
| $a_1(s) = \hat{a}_1 \left(\dfrac{W}{W_0} \exp\left[1 - \dfrac{W}{W_0}\right]\right)^{\alpha_1}$ | Attack rate on resource $R_1$ | m$^3$ day$^{-1}$ |
| $a_2(s) = \hat{a}_2 w^{\alpha_2}$ | Attack rate on resource $R_2$ | m$^2$ day$^{-1}$ |
| $H(s) = \chi_1 w^{\chi_2}$ | Handling time at size $s$ | day g$^{-1}$ |
| $p(s) = \left(\dfrac{p}{s}\right)^{\gamma_p} \left(s_0 - \dfrac{s_0}{1+\exp[-k_p(s-l_p)]}\right)$ | Predation mortality rate at size $s$ | day$^{-1}$ |
| $\mu_1(s) = m_1$ | Mortality in habitat 1 | day$^{-1}$ |
| $\mu_2(s) = m_2 + p(s)$ | Mortality in habitat 2 | day$^{-1}$ |
| $T_i(s,\mathbf{R}) = \dfrac{(a_i(s)R_i/\mu_i(s))^q}{(a_1(s)R_1/\mu_1(s))^q + (a_2(s)R_2/\mu_2(s))^q}$ | Fraction of time spent foraging in habitat $i$ | – |
| $I_i(s,\mathbf{R}) = \dfrac{a_i(s)R_i}{1+H(s)a_i(s)R_i}$ | Food intake rate from resource $R_i$ when feeding only on $R_i$ | g day$^{-1}$ |
| $f_i(s,\mathbf{R}) = T_i(s,\mathbf{R}) I_i(s,\mathbf{R})$ | Food intake rate from resource $R_i$ | g day$^{-1}$ |
| $f(s,\mathbf{R}) = f_1(s,\mathbf{R}) + f_2(s,\mathbf{R})$ | Total food intake rate | g day$^{-1}$ |
| $g(s,\mathbf{R}) = k(s) \dfrac{l_2 s}{(s/l_1)^{1/l_2}} \left(ef(s,\mathbf{R}) - b_1 w^{b_2}\right)$ | Growth rate | cm day$^{-1}$ |
| $b(s,\mathbf{R}) = \dfrac{q(1-k(s))}{h} \left(ef(s,\mathbf{R}) - b_1 w^{b_2}\right)$ | Birth rate | day$^{-1}$ |
| $d(s,\mathbf{R}) = T_1(s,\mathbf{R})\mu_1(s) + T_2(s,\mathbf{R})\mu_2(s)$ | Death rate | day$^{-1}$ |

**Supplementary Table 5.** Robustness of model results. A range of parameters was tested to determine the robustness of the conclusion that predation induces evolutionary divergence. The parameters are presented in descending order of sensitivity. Each parameter was tested for 10 or 11 equally spaced values for predation intensities 15%, 30%, 45%, 60%, and 75%, with gape size set to 18 cm.

| Parameter | Value | Tested range | Range producing divergence | Proportion of runs giving divergence |
|---|---|---|---|---|
| $\eta$ | 1 | [0.2,2] | [0.4,1.4] | 56 % |
| $w_0$ | 15 | [5,30] | [7.5,22.5] | 60 % |
| $q$ | 0.02 | [0.005,0.05] | [0.015,0.05] | 78 % |
| $\theta$ | 5 | [0,9] | [2,9] | 78 % |
| $m_1$ | 0.0025 | [0.0005,0.005] | [0.0005,0.004] | 78 % |
| $m_2$ | 0.0025 | [0.0005,0.005] | [0.0005,0.004] | 78 % |
| $\gamma_p$ | 1 | [0.5,1.5] | [0.6,1.5] | 90 % |
| $\kappa_a$ | 0.04 | [0.01,0.1] | [0.01,0.1] | 100 % |
| $k$ | 5 | [1,10] | [1,10] | 100 % |
| $k_p$ | 5 | [1,10] | [1,10] | 100 % |
